# The 14-3-3 Proteins Bmh1 and Bmh2 Are Key Regulators of Meiotic Commitment

**DOI:** 10.1101/2021.09.06.459189

**Authors:** Janardan N. Gavade, Chris Puccia, S. Grace Herod, Jonathan C. Trinidad, Luke E. Berchowitz, Soni Lacefield

**Affiliations:** Indiana University, Department of Biology, Bloomington, IN USA; Columbia University Irving Medical Center, Department of Genetics and Development, Hammer Health Sciences Center, New York, NY USA; Indiana University, Department of Chemistry, Bloomington, IN USA

**Keywords:** meiosis, meiotic commitment, return-to-growth (RTG), Cdc5, Polo kinase, Bmh1, Bmh2, 14-3-3 proteins, Ndt80, Ime2, Pes4

## Abstract

Induction of meiosis requires exogenous signals that activate internal gene regulatory networks. Meiotic commitment ensures the irreversible continuation of meiosis, even upon withdrawal of the meiosis inducing signals. Budding yeast cells have a unique property in that cells enter meiosis when starved, but if given nutrient-rich medium prior to the commitment point, they exit meiosis and enter mitosis. After the meiotic commitment point in prometaphase I, cells remain in meiosis even with addition of nutrients. Despite the importance of meiotic commitment in ensuring the production of gametes, only a few genes involved in the commitment process are known. We performed a genome-scale screen in budding yeast and discovered new regulators of meiotic commitment including Bcy1, which is involved in nutrient sensing, the meiosis-specific kinase Ime2, Polo kinase Cdc5, and the 14-3-3 proteins Bmh1 and Bmh2. Importantly, we found that Bmh1 and Bmh2 are involved in multiple processes throughout meiosis including the maintenance of the middle meiosis transcription factor Ndt80, activation of Cdc5, and interaction with an RNA-binding protein Pes4, which is important for regulating the timing of translation of several mRNAs in meiosis II. This study identifies a meiotic commitment regulatory network with the 14-3-3 proteins functioning as central regulators.

## Introduction

The production of haploid gametes requires the cell division process of meiosis, in which one round of DNA replication is followed by two rounds of chromosome segregation. Meiotic entry occurs in the presence of extrinsic signals that activate intrinsic gene regulatory networks (Kimble, 2011; van Werven & Amon, 2011). Precise control of meiotic regulators ensures that cell cycle events progress in a particular order (MacKenzie & Lacefield, 2020). Some of the pathways that promote mitotic cell division are inhibited during some meiotic stages to ensure that cells complete meiosis (Okaz et al., 2012; Wei et al., 2014; Weidberg, Moretto, Spedale, Amon, & van Werven, 2016).

The importance of inhibiting mitotic cell-cycle regulators during meiosis is demonstrated through the identification of genetic mutants that form germ cell tumors due to the misexpression of mitotic regulators during meiosis. For example, in *C. elegans* and *D. melanogaster*, mutations in the translation repressors GLD-1 and Bruno, respectively, cause inappropriate expression of mitotic cyclins during oogenesis. Once expressed, the mitotic cyclins bound and activated cyclin-dependent kinase (CDK) and the cells entered mitosis, which caused germ cell tumor formation (Biedermann et al., 2009; Sugimura & Lilly, 2006).

Although these studies have revealed important aspects of meiotic maintenance, we have only a limited understanding of the factors that allow cells to complete meiosis by preventing the transition to mitosis. To address the question of how cells maintain meiosis, we chose to study meiotic commitment in *S. cerevisiae.* We define meiotic commitment as the point at which the meiosis-inducing signal is no longer required for cells to complete meiosis (Tsuchiya, Yang, & Lacefield, 2014; Winter, 2012). In budding yeast, starvation is the major meiosis-inducing signal and cells need to remain starved until reaching the commitment point in mid-prometaphase I. Although many organisms have key transition points before the entrance into the meiotic divisions, budding yeast is unique in its natural ability to return to mitosis if the meiosis-inducing signal is not maintained prior to the meiotic commitment point (Esposito & Esposito, 1974; Kimble, 2011; Sherman & Roman, 1963; Simchen, 2009; Simchen, Pinon, & Salts, 1972; Winter, 2012). This unique cell cycle, transitioning from meiosis to mitosis, is termed ‘return-to-growth’. If nutrient-rich medium is provided after the commitment point, cells remain in meiosis, complete meiosis, and package the four meiotic products into spores.

Maintaining meiotic commitment is important for genomic integrity. In meiotic commitment mutants, upon the addition of nutrient-rich medium, cells inappropriately exit meiosis after the first nuclear division creating multi-nucleate polyploid cells after the subsequent mitotic division (Ballew & Lacefield, 2019a, 2019b; Tsuchiya et al., 2014). High levels of the Ndt80 transcription factor are important for maintaining meiotic commitment (Tsuchiya et al., 2014). Ndt80 induces the transcription of middle meiosis genes whose protein products are needed for prophase I exit and the meiotic divisions (Chu et al., 1998; Chu & Herskowitz, 1998; Hepworth, Friesen, & Segall, 1998; Winter, 2012; Xu, Ajimura, Padmore, Klein, & Kleckner, 1995).

To identify new regulators of meiotic commitment, we performed a genome-scale screen. Because high Ndt80 levels are needed for meiotic commitment, we used strain with reduced levels of Ndt80 and screened for genes that when overexpressed rescued meiotic commitment. We identified five proteins that promote meiotic commitment: Bcy1, a regulator of nutrient sensing; Ime2, a CDK-like kinase that activates Ndt80 to maintain meiotic commitment; Polo kinase, Cdc5, that regulates several steps in meiosis; and 14-3-3 proteins Bmh1 and Bmh2. Importantly, we found that Bmh1 and Bmh2 are involved in multiple processes throughout meiosis that affect meiotic commitment. Bmh1 and Bmh2 are needed for normal Ndt80 levels, activation of Polo kinase, and interaction with an RNA-binding protein, Pes4, which regulates the timing of translation of several mRNAs important for meiosis II progression and gamete formation. Our study supports a model where meiotic commitment is actively maintained through multiple processes throughout meiosis.

## Results

### *BCY1, IME2, BMH1, BMH2*, and *CDC5* are important for meiotic commitment

We monitor commitment using a microfluidics setup, which allows us to flow in nutrients at precise stages of meiosis. We use fluorescent markers to determine the meiotic stage at the time of nutrient addition. (Carminati & Stearns, 1997; Scherthan et al., 2007; Tsuchiya et al., 2014). Zip1-GFP serves as a marker for prophase I, as the synaptonemal complex is assembled and disassembled in prophase I (Scherthan et al., 2007; Sym, Engebrecht, & Roeder, 1993; White, Cowan, Cande, & Kaback, 2004). Tub1-GFP marks the spindle and Spc42-mCherry marks the spindle pole bodies (SPBs), which allow us to determine the stages beyond prophase I (Tsuchiya & Lacefield, 2013). Although Zip1 and Tub1 are both tagged with GFP, they are easy to distinguish due to the temporal and morphological differences of the synaptonemal complex and spindle. If nutrients are added to cells in early prometaphase I or earlier stages, the cells exit meiosis, form a bud, and undergo mitosis (Fig. 1A-E)(Tsuchiya et al., 2014). Cells in late prometaphase I and stages beyond are committed to meiosis and will stay in meiosis upon nutrient addition (Figure 1B,D,E).

**Figure 1:**
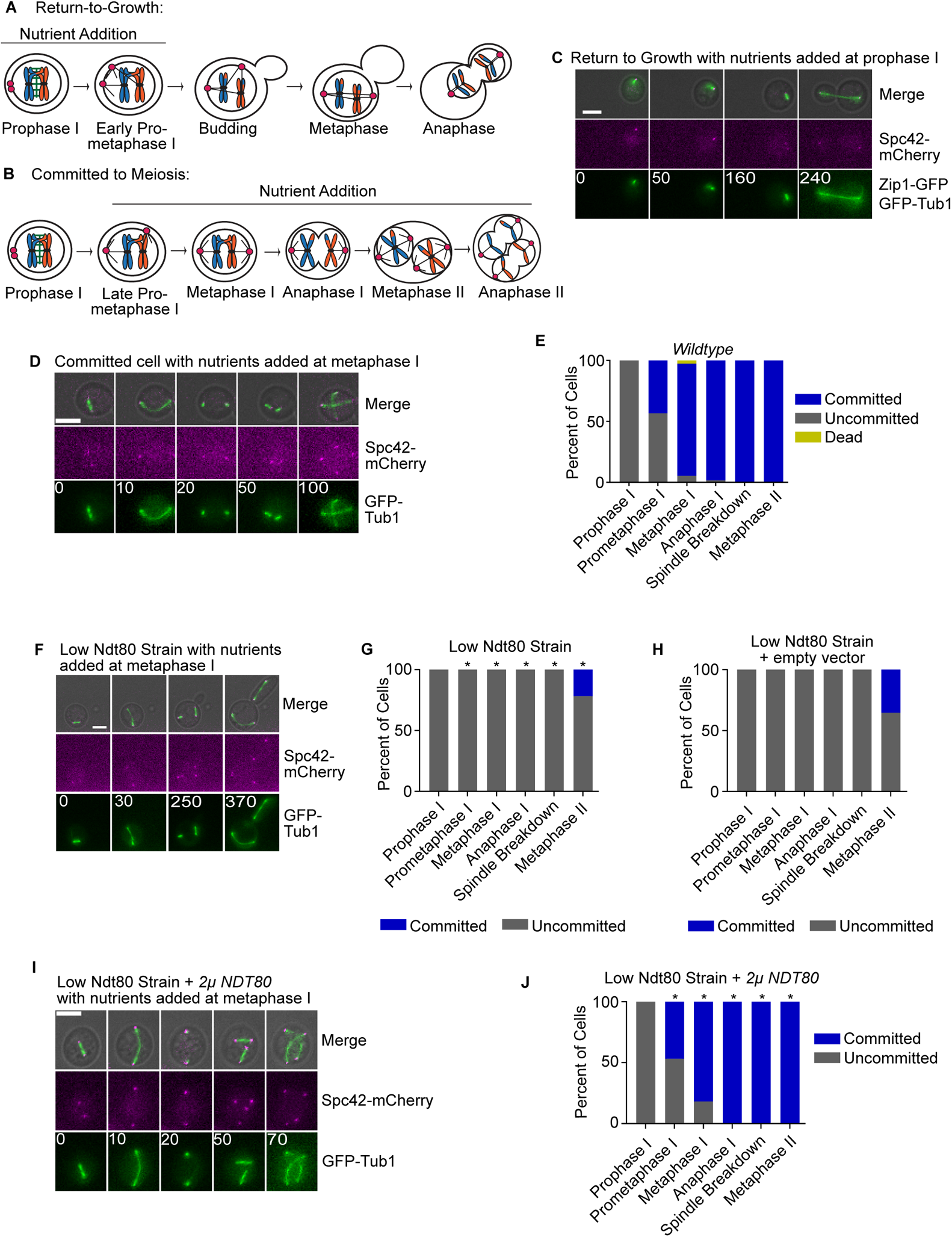
A genetic screen to identify the genes involved in meiotic commitment. (A) Cartoon of a cell undergoing Return-to-Growth (RTG) upon nutrient addition at prophase I/ early prometaphase I. (B) Cartoon of a cell committed to meiosis commitment upon nutrient addition in late prometaphase I to anaphase II. (C) Time-lapse image of a wildtype cell undergoing RTG upon nutrient addition in pachytene (t = 0). Time in minutes. Scale bar −5µm. (D) Time-lapse images of a wildtype cell committed to meiosis upon nutrient addition at metaphase I (t = 0). Time in minutes. Scale bar – 5µm. (E) Graph of the commitment assay for a wildtype strain with 268 cells counted over two independent experiments. (F) Time-lapse images of a low Ndt80 cell with nutrient addition at metaphase I (t = 0). Time in minutes. Scale bar – 5µm. (G) Graph of the commitment assay for the low Ndt80 strain. 199 cells were counted in two independent assays. This mutant displayed a statistically significant difference from wildtype at prometaphase I - metaphase II (*p<0.05, Fisher’s exact test). (H) Graph of the commitment assay for the low Ndt80 strain with 2µ empty vector. 256 cells counted in 2 independent experiments. The x-axis depicts the stage at the time of nutrient addition. This mutant is not significantly different from the low Ndt80 strain without an empty vector (Fisher’s exact test). (I) Time-lapse images of low Ndt80 cell with a 2µ-NDT80 vector with nutrient addition at metaphase I (t = 0). Time in minutes. Scale bar – 5µm. (J) Graph of the commitment assay for the low Ndt80 + 2µ-NDT80 vector strain. 136 cells counted in two independent experiments. The X-axis depicts the stage at the time of nutrient addition. This mutant displayed a statistically significant difference from Low Ndt80 cells in prometaphase I - metaphase II stages (*p<0.05, Fisher’s exact test).

Reducing *NDT80* transcription by deleting the mid-sporulation elements within the promoter results in low levels of Ndt80 and ablates meiotic commitment (Tsuchiya et al., 2014). The cells returned to mitosis if nutrients were added at any stage of meiosis, often causing multi-nucleate polyploid cells (Fig. 1F-G)(Tsuchiya et al., 2014). For ease of discussion, we will call this strain the “low Ndt80 strain” throughout the manuscript. The low Ndt80 strain likely fails to commit to meiosis due to the low expression of an Ndt80 target gene. Therefore, we performed a systematic screen to identify genes that when overexpressed rescue meiotic commitment (Fig. S1A,B). We allowed the low Ndt80 cells with an individual 2μ plasmid from the Yeast Tiling Collection to reach meiosis II, added nutrients, and then imaged the cells to score as either uncommitted or committed (G. M. Jones et al., 2008). Uncommitted cells, such as those with the empty vector control, budded and underwent mitosis, even when nutrients were added to cells at metaphase I and beyond (Fig. S1B, 1H). Committed cells either remained in meiosis after nutrient addition in stages beyond metaphase I or completed meiosis (Fig. S1B). *NDT80* on a 2μ plasmid rescued meiotic commitment and served as a positive control (Fig. 1I,J).

We confirmed our positive hits and focused on the plasmids identified in both replicates (Table S1). Because the plasmids contain multiple genes, we subcloned individual genes of interest on a 2μ plasmid and tested if they suppressed the meiotic commitment defects in the low Ndt80 strains. We chose to focus on five genes that suppressed the loss of commitment: *BCY1, IME2*, *BMH1*, *BMH2*, and *CDC5*. The identification of *BCY1* and *IME2* in our genetic screen was not surprising due to their known functions in inhibiting the glucose response pathway and activating Ndt80, respectively (Benjamin, Zhang, Shokat, & Herskowitz, 2003; Broach, 2012; Sopko, Raithatha, & Stuart, 2002). In contrast, *BMH1*, *BMH2*, and *CDC5* have known roles in meiosis, but they do not have known functions that would implicate a role in meiotic commitment. We focused on characterization of these genes to possibly reveal a pathway for maintaining meiotic commitment.

### Bcy1 and Ime2 ensure meiotic commitment

Bcy1 is a negative regulatory subunit of protein kinase A (PKA), which is part of the RAS/cAMP-dependent signaling pathway induced in response to glucose (Broach, 2012; Thevelein & de Winde, 1999; Toda et al., 1987). We found that overexpression of *BCY1* increased the population of committed cells when nutrient-rich medium is added to the low Ndt80 strain in metaphase I, anaphase I, and metaphase II (Fig. S1C). Conversely, reducing the levels of Bcy1 by deleting one copy of *BCY1* in a wildtype strain resulted in an increased population of uncommitted cells when compared to wildtype (Fig. S1D, 1E, p<0.05, Fisher’s exact test). We chose to reduce the levels, instead of making a homozygous deletion, because some Bcy1 is needed for cells to enter meiosis (Matsuura et al., 1990; Tripp & Pinon, 1986). Overall, these results demonstrate that full levels of Bcy1 are needed for meiotic commitment, likely by preventing the activation of PKA.

Ime2 is a meiosis-specific kinase that has a known role in activating Ndt80 through phosphorylation (Benjamin et al., 2003; Sopko et al., 2002). Therefore, it was not surprising that increased expression of *IME2* resulted in a greater population of committed cells when compared to the low Ndt80 strain without overexpression of *IME2* (Fig. S1E,1E; p<0.05 Fisher’s exact test). We predicted that decreased Ime2 activity would result in a loss of commitment in an otherwise wildtype strain. We mutated two residues in the Ime2 activation loop, T242A and Y244F. Previous studies revealed that these residues are phosphorylated for full Ime2 activity (Schindler et al., 2003; Schindler & Winter, 2006). Therefore, we asked if meiotic commitment was disrupted in the point mutants. Because Ime2 also phosphorylates the *NDT80* repressor Sum1 to help release Sum1 from the *NDT80* promoter, we performed these experiments in *sum1Δ* cells (Ahmed, Bungard, Shin, Moore, & Winter, 2009; Lo et al., 2012; Moore et al., 2007; Pak & Segall, 2002). Loss of *SUM1* did not affect meiotic commitment (Fig. S1F). In contrast, the *ime2-T242A sum1Δ* mutants had a strong defect in meiotic commitment in that all metaphase I and anaphase I cells were uncommitted and exited meiosis upon nutrient-rich medium addition (Fig. S1G). The *ime2-Y244F sum1Δ* mutant did not enter the meiotic divisions in our strain background, preventing us from analyzing meiotic commitment. Overall, our results support the model that full Ime2 activity is required for meiotic commitment.

### Bmh1 and Bmh2 Are Required for Full Ndt80 Expression and Commitment

We next focused on *BMH1* and *BMH2*, which encode the two 14-3-3 proteins in budding yeast (Aitken et al., 1992; van Hemert, van Heusden, & Steensma, 2001; van Heusden, Wenzel, Lagendijk, de Steensma, & van den Berg, 1992). 14-3-3 proteins typically bind phosphorylated residues on proteins to modulate their activity, stimulate protein-protein interactions, regulate sub-cellular localization, affect their stability, and even facilitate chaperone-like activities (Braselmann & McCormick, 1995; H. K. Chen et al., 2003; Lopez-Girona, Furnari, Mondesert, & Russell, 1999; Muslin, Tanner, Allen, & Shaw, 1996; Obsil, Ghirlando, Klein, Ganguly, & Dyda, 2001; Sluchanko & Gusev, 2017; Yaffe et al., 1997). In the low Ndt80 strain, overexpression of *BMH1* or *BMH2* resulted in a higher population of committed cells as compared to the low *NDT80* strain with the empty vector (Fig. 2A-C, 1H; p<0.05 Fisher’s exact test). We scored cells as committed if they either completed meiosis or arrested in meiosis I or meiosis II (Fig. 2C).

**Figure 2:**
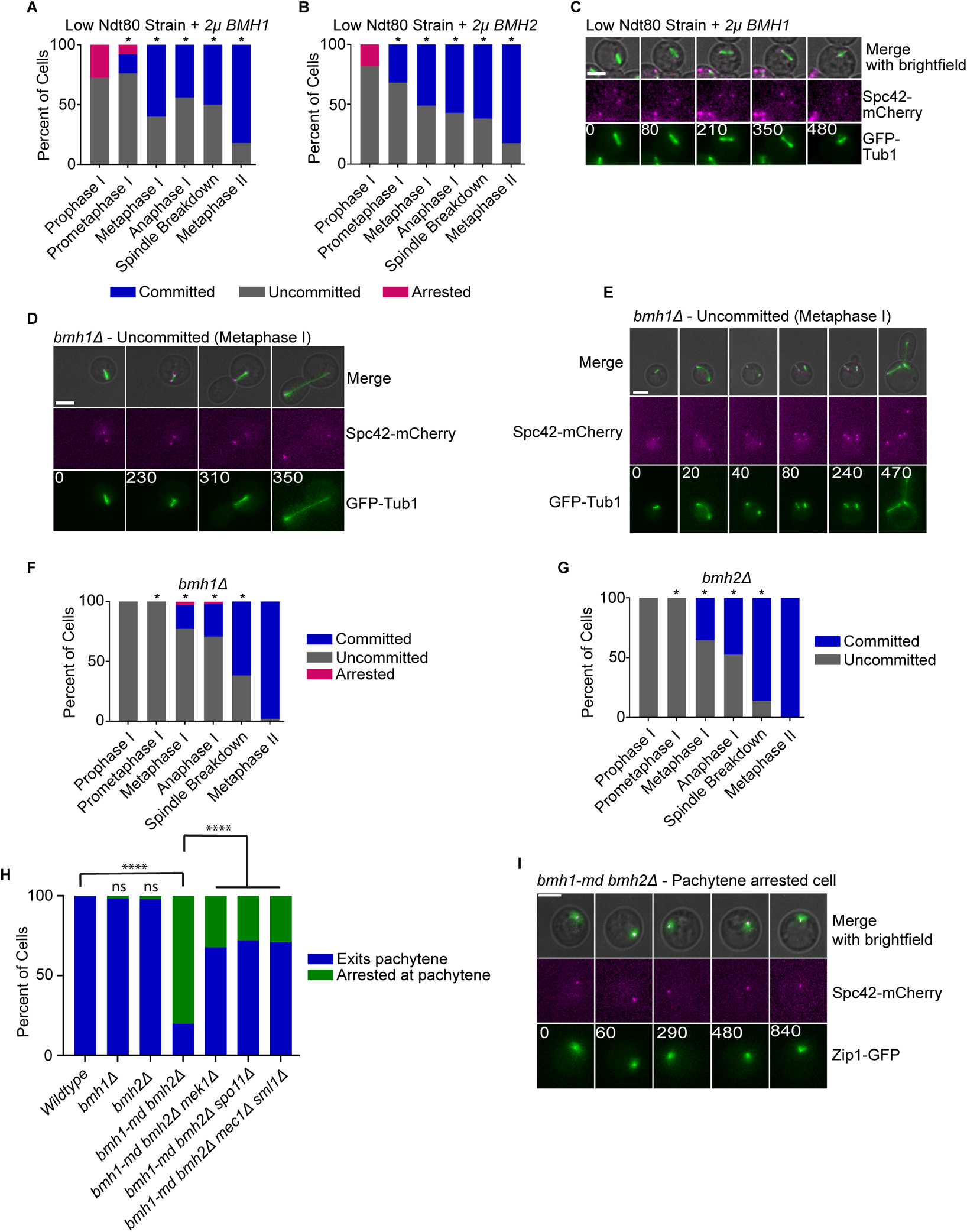
Bmh1 are Bmh2 are important for meiotic commitment and passage through the pachytene checkpoint. (A) Graph of the commitment assay for the low Ndt80 strain with 2µ-*BMH1*. 243 cells were counted in two independent experiments. The x-axis shows the stage at the time of nutrient addition. This mutant displayed a statistically significant difference from the low Ndt80 strain in prometaphase I - metaphase II (*p<0.05, Fisher’s exact test). (B) Graph of the commitment assay for the low Ndt80 strain with 2µ-*BMH2*. 227 cells from two independent experiments were counted. The x-axis shows the stage at the time of nutrient addition. This mutant displayed a statistically significant difference from the low Ndt80 strain in prometaphase I - metaphase II (*p<0.05, Fisher’s exact test). (C) Time-lapse images of a committed low Ndt80 + 2µ-*BMH1* cell with nutrient addition at metaphase I (t = 0). Time in minutes. Scale bar – 5µm. (D and E) Time- lapse images of *bmh1Δ* uncommitted cells upon nutrient addition in metaphase I (t = 0). Time in minutes. Scale bar - 5µm. (F) Graph of the commitment assay for the *bmh1Δ* strain. 288 cells were counted over three independent experiments. The x-axis shows the stage at the time of nutrient addition. This mutant displayed a statistically significant difference from wildtype cells in prometaphase I - spindle breakdown (*p<0.05, Fisher’s exact test). (G) Graph of the commitment assay for the *bmh2Δ* strain. 363 cells were counted over three independent experiments. The x-axis shows the stage at the time of nutrient addition. This mutant displayed a statistically significant difference from the wildtype strain in prometaphase I - spindle breakdown stages (*p<0.05, Fisher’s exact test). (H) Graph of the progression through pachytene for the indicated strains. 300 cells were counted per genotype from three independent experiments for each genotype. Asterisks indicate statistically significant difference (**** p<0.0001, Fisher’s exact test), ns is not significant. (I) Time-lapse images of a *bmh1-md bmh2Δ* double mutant cell arrested at pachytene for 14 hours. Time zero is 8 hrs after SPM transfer. Time in minutes. Scale bar - 5µm.

To determine if 14-3-3 proteins are required for meiotic commitment, we analyzed single deletions of *BMH1* and *BMH2* in otherwise wildtype strains. Previous reports showed that *bmh1Δ* and *bmh2Δ* mutants exhibit minor sporulation and spore viability defects (Kumar, 2018; Slubowski, Paulissen, & Huang, 2014). We found that both *bmh1Δ* and *bmh2Δ* strains were severely defective for meiotic commitment when nutrients were added in metaphase I and anaphase I (Fig. 2D-G). We observed two phenotypes of the uncommitted cells that had nutrients added in prometaphase I or metaphase I: i) cells formed a bud and underwent a mitotic division (Fig. 2D); or, ii) cells underwent meiosis I, formed a bud and then underwent a mitotic division with two spindles (Fig. 2E).

The C-terminus of 14-3-3 proteins interacts with target proteins and the phosphorylation of a conserved residue within the C-terminus is important for efficient binding (Ichimura et al., 1995; Liu, Elly, Yoshida, Bonnefoy-Berard, & Altman, 1996; Luo, Zhang, Rapp, & Avruch, 1995; Wakui, Wright, Gustafsson, & Zilliacus, 1997). In Bmh1, the S235 residue resides in a patch of amino acids that are highly conserved among 14-3-3 orthologous proteins in fungi, plants, worms, zebrafish, flies, mouse, and humans (Fig. S2A). To determine if Bmh1 phosphorylation was likely important for meiotic commitment, we mutated S235 to alanine as well as S238 and S240, due to their proximity, making a triple mutant (*bmh1-3A*). We found that *bmh1-3A* cells had an increased population of uncommitted cells compared to wildtype when nutrients are added in metaphase I and anaphase I (Fig.S2B, 1E; p<0.05 Fisher’s exact test). Therefore, the serine residues within the Bmh1 C-terminus are required for proper meiotic commitment.

*BMH1* and *BMH2* are paralogs with redundant functions (van Hemert, van Heusden, et al., 2001). They can bind their substrates as monomers, homodimers, and as heterodimers (Chaudhri, Scarabel, & Aitken, 2003; Fischer et al., 2009; D. H. Jones, Ley, & Aitken, 1995). But, Bmh1 and Bmh2 mostly form heterodimers (Chaudhri et al., 2003). We hypothesized that some cells remained committed in the single mutants due to the presence of the other paralog. In our strain background, the double knockout mutant is not viable. Therefore, to make a double mutant in meiosis, we made a meiotic depletion allele by placing *BMH1* under the control of the mitosis-specific *CLB2* promoter and we deleted *BMH2* (*bmh1-md bmh2Δ*). We found that 80% of the cells arrested at the pachytene stage of prophase I, scored by Zip1-GFP presence (Fig. 2H,I). Because the *bmh1-md bmh2Δ* cells enter prophase I, unlike the double knockouts reported in a different strain background (Kumar, 2018), we assume that there is a low level of Bmh1 protein present. The 20% of *bmh1-md bmh2Δ* cells that exited prophase I then arrested at several different stages of meiosis, demonstrating that Bmh1 and Bmh2 have important functions throughout meiosis (Fig. S2C).

We asked whether the *bmh1-md bmh2Δ* cells that arrested in pachytene triggered the pachytene checkpoint, which monitors DNA damage. We deleted two genes important for signaling the checkpoint, *MEK1* and *MEC1*. Mek1is a meiosis-specific checkpoint kinase downstream of the checkpoint signal (X. Chen et al., 2018; Hollingsworth & Gaglione, 2019; Prugar, Burnett, Chen, & Hollingsworth, 2017; Rockmill & Roeder, 1991). Mec1 responds to DNA damage and transduces that signal for a checkpoint arrest (Lydall, Nikolsky, Bishop, & Weinert, 1996; Subramanian & Hochwagen, 2014; Weinert, Kiser, & Hartwell, 1994). *MEC1* is an essential gene, but *mec1Δ* cells can survive with a deletion of *SML1* (Zhao, Muller, & Rothstein, 1998). The majority of *bmh1-md bmh2Δ mek1Δ* and *bmh1-md bmh2Δ mec1Δ sml1Δ* cells exit prophase I, suggesting that the cells were arrested due to the activation of the pachytene checkpoint (Fig. 2H). To determine if the pachytene checkpoint arrest was due to unrepaired programmed double strand breaks (DSBs), we deleted the *SPO11* gene, which encodes the enzyme that makes programmed DSBs during meiosis (Keeney, Giroux, & Kleckner, 1997). We find that the majority of *spo11Δ bmh1-md bmh2Δ* cells also exit prophase I (Fig. 2H). These results suggest that Bmh1 and Bmh2 likely have important roles in the repair of programmed DSBs. Although this function is intriguing, we decided to focus on the roles of Bmh1 and Bmh2 in meiotic commitment.

We considered that Ndt80 could directly interact with Bmh1 and Bmh2. Using a two-hybrid assay, we identified an interaction between *BMH1* and a fragment of *NDT80* (amino acids 287-627), which does not contain the DNA-binding domain (Fig. 3A). Therefore, Bmh1 could directly regulate Ndt80 activity or localization by binding Ndt80.

**Figure 3:**
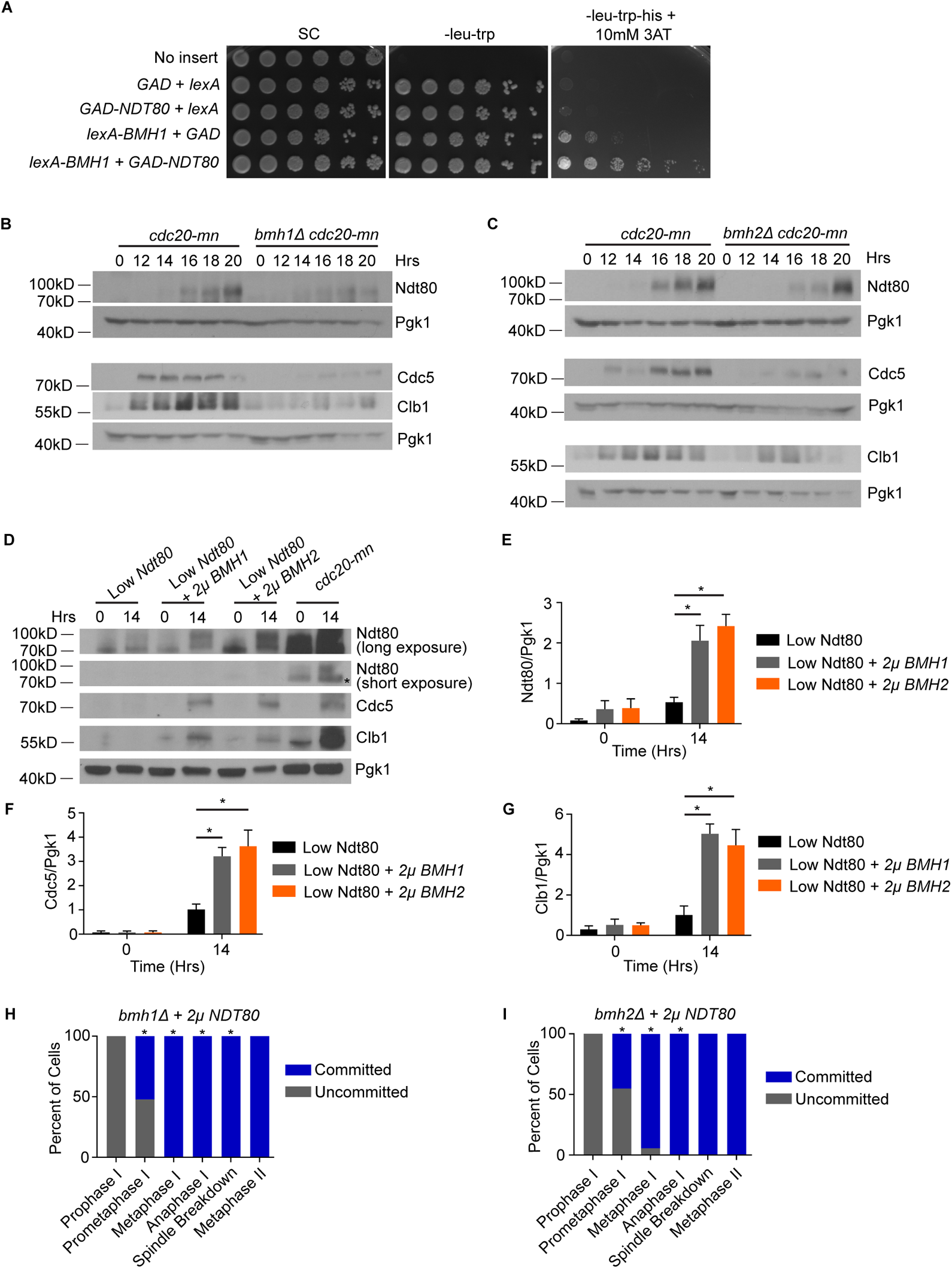
Bmh1 and Bmh2 regulate Ndt80 levels. (A) Yeast 2-hybrid were spotted on plates using a 10-fold serial dilution. (B and C) Western blot of Ndt80 levels, Cdc5 levels, and Clb1 levels in *cdc20mn*, *bmh1Δ cdc20mn* (B), and *cdc20mn, bmh2Δ cdc20mn* cells (C). Time zero indicates the time at which cells were transferred to SPM medium. Pgk1 serves as the loading control. (D) Western blotting analysis of Ndt80 levels, Cdc5 levels, and Clb1 levels in Low Ndt80, Low Ndt80 +2µ*-BMH1*, Low Ndt80 + 2µ*-BMH2,* and *cdc20-mn* cells. Time zero indicates the time at which cells were transferred into SPM medium. Pgk1 is the loading control. (E-G) The graphs depict the mean ± SEM of the densitometric analysis of Ndt80 (E), Cdc5 (F) and Clb1 (G) levels relative to Pgk1 in the indicated strains at indicated time points. The difference in the Ndt80/Pgk1, Cdc5/Pgk1 and Clb1/Pgk1 levels is statistically significantly different at the t = 14 timepoint in the low Ndt80 +2µ*-BMH1* and low Ndt80 + 2µ*-BMH2* compared to the Low Ndt80 strain (* indicates p<0.05, Unpaired t-test with Welch’s correction). (H) Graph of the commitment assay for *bmh1Δ* + 2µ-*NDT80*. 233 cells were counted in two independent experiments. The x-axis shows the stage at the time of nutrient addition. This mutant displayed a statistically significant difference from *bmh1Δ* cells in prometaphase I - spindle breakdown (* p<0.05, Fisher’s exact test). (I) Graph of the commitment assay for *bmh2Δ* + 2µ-*NDT80* strain. 246 cells were counted in two independent experiments. The x-axis depicts the stage at the time of nutrient addition. This mutant displayed a statistically significant difference from *bmh2Δ* cells in prometaphase I - anaphase I (* p<0.05, Fisher’s exact test)

Because Ndt80 increases its own transcription, a decrease in Ndt80 activity or localization could cause a decrease in Ndt80 protein levels (Hepworth et al., 1998; Tsuchiya et al., 2014; Tung, Hong, & Roeder, 2000; Winter, 2012). Therefore, we measured the levels of Ndt80 in *bmh1Δ* and *bmh2Δ* single mutants arrested at metaphase I (due to a Cdc20 meiotic null). We found that Ndt80 levels were strongly reduced in *bmh1Δ cdc20-mn* and *bmh2Δ cdc20- mn* cells when compared to *cdc20-mn* cells (Fig. 3B-C, quantification in Fig. S3A-B; p<0.05, Two-way Anova). Furthermore, the protein levels of Clb1 and Cdc5, whose production is dependent on Ndt80, were reduced in *bmh1Δ cdc20-mn* and *bmh2Δ cdc20-mn* cells compared to *cdc20-mn* cells (Fig. 3B-C, quantification in Fig. S3C-F; p<0.05, Two-way Anova).

Because 14-3-3 proteins can modulate the activity of the proteins that they bind, we asked if the overexpression of Bmh1 and Bmh2 resulted in increased levels of Ndt80 or Ndt80 target proteins in the low Ndt80 strains (Obsil et al., 2001). We compared Ndt80 levels in low Ndt80 strains with or without a *BMH1* or *BMH2* 2μ plasmid. We find that the levels of Ndt80 are increased with *BMH1* and *BMH2* overexpression (Fig. 3D-E; p<0.05, Unpaired t-test with Welch’s correction). The levels of the Ndt80 targets Cdc5 and Clb1 are also increased compared to the low Ndt80 strain (Fig. 3D, F-G; p<0.05, Unpaired t-test with Welch’s correction). These results suggest that *BMH1* and *BMH2* overexpression rescued the commitment defect of the low Ndt80 strain by increasing the levels of Ndt80 and Ndt80-dependent targets. To test this idea, we asked whether overexpression of *NDT80* rescues meiotic commitment in *bmh1Δ* and *bmh2Δ* cells. We transformed *bmh1Δ* and *bmh2Δ* cells with a 2μ plasmid containing *NDT80*. We found that the increased expression of *NDT80* rescued meiotic commitment in *bmh1Δ* and *bmh2Δ* cells (Fig. 3H,I). Overall, these results reveal that Bmh1 and Bmh2 maintain the levels of Ndt80 that are required for proper meiotic commitment.

### Bmh1 Interacts with RNA-binding protein Pes4 to maintain meiotic commitment

Although reduced Ndt80 levels in *bmh1Δ* and *bmh2Δ* mutants could explain the commitment defect, we hypothesized that Bmh1 and Bmh2 may interact with other proteins required for meiotic commitment. To identify other proteins, we immunoprecipitated Bmh1 in meiosis and assessed the bound proteins by mass spectrometry. For the immunoprecipitation, we tagged the C-terminus of Bmh1 with GFP and found that addition of the tag did not disrupt Bmh1 function, as assayed by survival and normal growth in a *bmh2Δ* background. We compared the proteins immunoprecipitated from the strain with Bmh1-GFP to an untagged control strain and only considered those isolated from Bmh1-GFP. We found many interesting interacting proteins, including an RNA-binding protein called Pes4 (Suppl. Table S2). *PES4* transcription is induced by Ndt80 (Chu et al., 1998). Although Pes4 is not required for meiosis, the known function of Pes4 is to delay the translation of a subset of Ndt80-dependent mRNAs until the end of meiosis II (Jin, Zhang, Sternglanz, & Neiman, 2017). Interestingly, a previous study showed that with nutrient-rich medium addition after the meiotic commitment point, most Ndt80-dependent mRNAs are unstable, except for a handful of mRNAs that are protected (Friedlander et al., 2006). The protection of some of these mRNAs was shown to depend on Pes4 (Jin et al., 2017).

To determine if Pes4 had an important role in meiotic commitment, we performed commitment assays in *pes4Δ* cells. We found that a significant increase in the number of uncommitted cells in *pes4Δ* cells compared to wildtype cells when nutrient-rich medium was added in metaphase I and anaphase I (Fig. 4A, 1E; p<0.05 Fisher’s exact test). Furthermore, overexpression of Pes4 in *bmh1Δ* and *bmh2Δ* rescued meiotic commitment in prometaphase I, metaphase I, anaphase I, and metaphase II when compared to the single mutants without *PES4* overexpression (Fig. 4B, 2F,G; p<0.05 Fisher’s exact test). In summary, our results support the idea that Bmh1interacts with Pes4 to ensure meiotic commitment.

**Figure 4:**
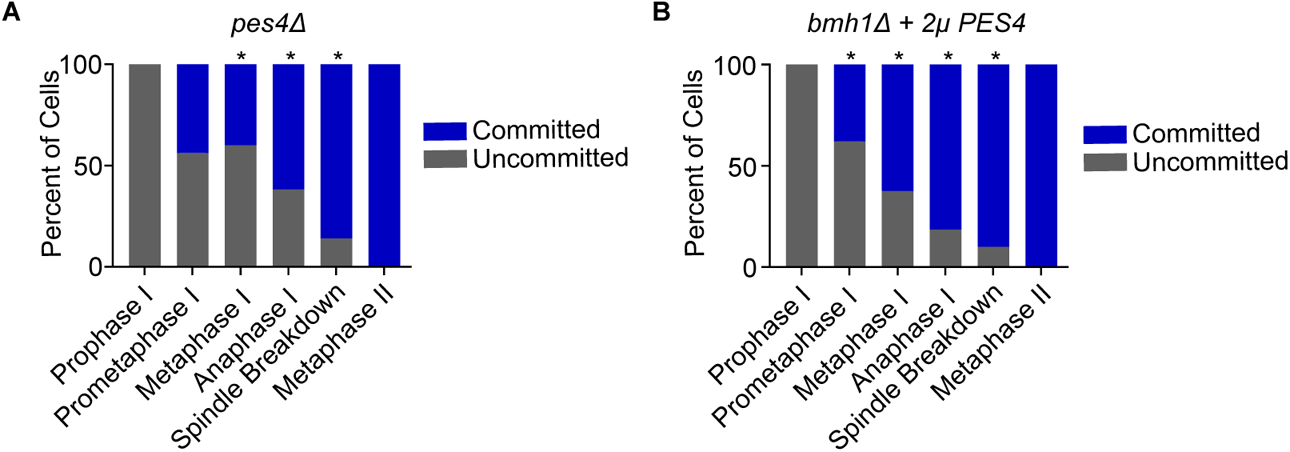
Pes4 is required for meiotic commitment and *PES4* overexpression rescues commitment in *bmh1Δ* mutants. (A) Graph of the commitment assay for the *pes4Δ* strain. 186 cells in two independent experiments were counted. The x-axis shows the stage at the time of nutrient addition. This mutant displayed a statistically significant difference from wildtype cells in metaphase I, anaphase I, and spindle breakdown stages (*p<0.05, Fisher’s exact test). (B) Graph of the commitment assay for the *bmh1Δ +* 2µ*-PES4* strain. 181 cells were counted in two independent experiments. This mutant displayed a statistically significant difference from *bmh1Δ* cells in prometaphase I - spindle breakdown (*p<0.05, Fisher’s exact test).

### Polo Kinase is Essential for Meiotic Commitment

Although we uncovered *CDC5* in our genetic screen, overexpression of *CDC5* only modestly rescued the low Ndt80 strain (Fig. 5A). *CDC5* expression is induced by Ndt80 and although Cdc5 phosphorylates Ndt80, it was shown not to regulate the levels of the Ndt80 target genes tested (Chu et al., 1998; Clyne et al., 2003). These prior findings suggested that Cdc5 does not regulate meiotic commitment through Ndt80. Cdc5 has several important roles in meiosis I (Clyne et al., 2003; B. H. Lee & Amon, 2003; Sourirajan & Lichten, 2008). We hypothesized that Cdc5 may be important for meiotic commitment, but that the overexpression does not fully rescue the low Ndt80 strain because other proteins downstream of Cdc5 remain limiting. To test this hypothesis, we made a meiotic null allele of Cdc5 by placing *CDC5* under the control of the mitosis-specific *CLB2* promoter (*cdc5-mn*). In the absence of Cdc5, cells arrest in a metaphase I-like state, with a short bipolar spindle (Clyne et al., 2003; B. H. Lee & Amon, 2003). We compared the *cdc5-mn* cells to *cdc20-mn* cells that also arrest at metaphase I (due to a lack of APC/C activity). However, because Cdc5 is important for synaptonemal complex disassembly and the clamping of sister chromatid kinetochores, the *cdc5-mn* cells arrest with synaptonemal complex and with sister chromatid kinetochores bioriented instead of homologous chromosome kinetochores (Clyne et al., 2003; B. H. Lee & Amon, 2003; Sourirajan & Lichten, 2008)(Fig. 5B). With nutrient addition, all *cdc20-mn* cells are committed and remained arrested at metaphase I (Fig. 5C,D). In contrast, 85% of *cdc5-mn* cells exit meiosis, form a bud, and undergo a mitotic division (Fig. 5D,E). These results suggest that Cdc5 is required for meiotic commitment.

**Figure 5:**
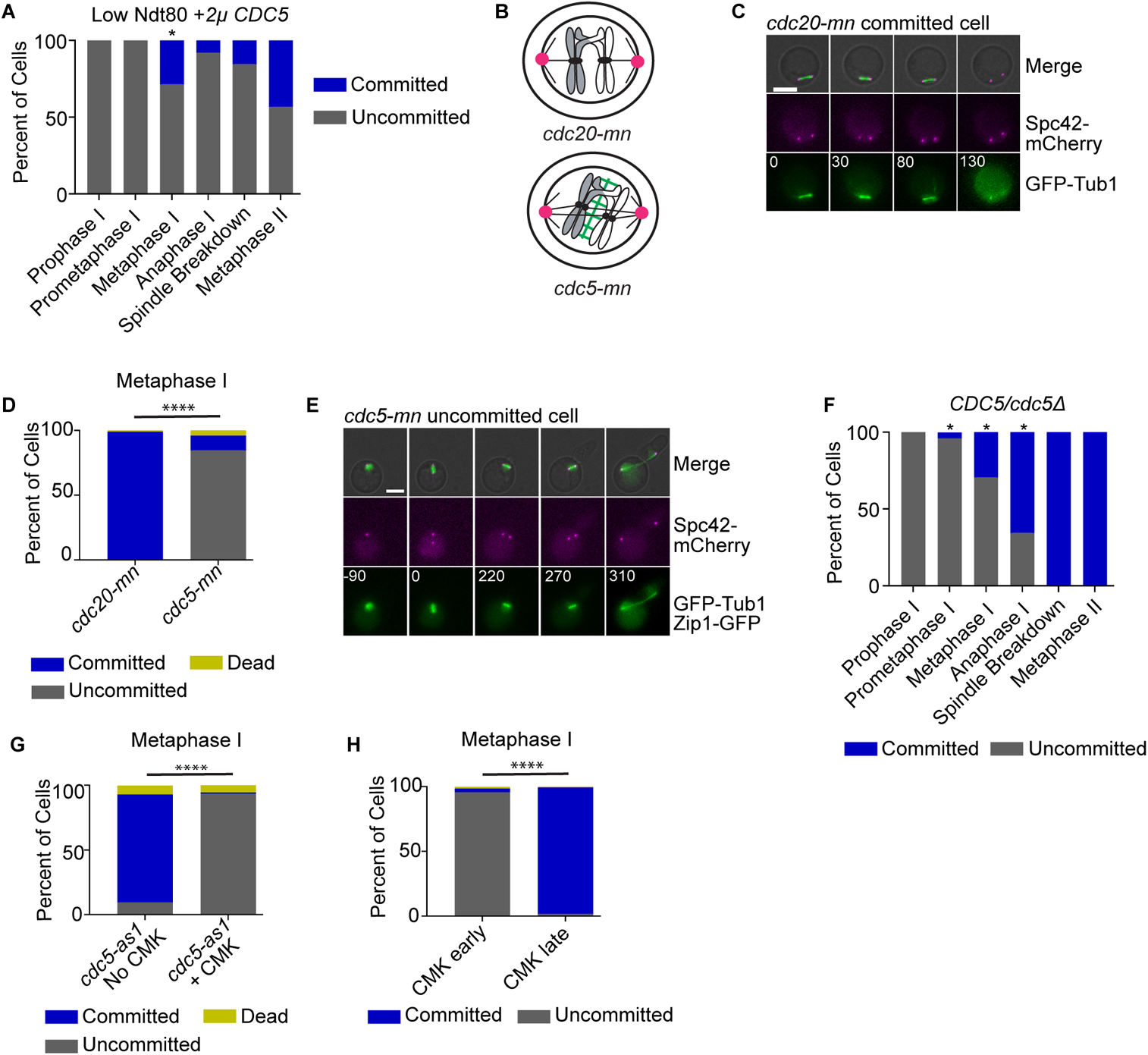
Cdc5 is important for meiotic commitment. (A) Graph of the commitment assay for the low Ndt80 +2µ-*CDC5* strain. 202 cells were counted in two independent experiments. This strain displayed a statistically significant difference from the low Ndt80 strain in metaphase I (*p<0.05, Fisher’s exact test). (B) Cartoon of chromosomal biorientation in *cdc20-mn* and *cdc5-mn* cells. (C) Time-lapse images of a *cdc20-mn* committed cell with nutrient addition at metaphase I (t = 0). Time in minutes. Scale bar - 5µm. (D) Graph of the commitment assay for the *cdc20-mn* strain (n = 113 cells; two independent experiments) and *cdc5-mn* strain (n = 156; three independent experiments). All cells were scored from nutrient addition at metaphase I. Asterisks indicate statistically significant difference (**** p<0.0001, Fisher’s exact test). (E) Time-lapse images of a *cdc5-mn* uncommitted cell with nutrient addition from metaphase I (t = 0). Time in minutes. Scale bar - 5µm. (F) Graph of the commitment assay for *CDC5/cdc5Δ* heterozygous strain. 207 cells from two independent experiments were counted. The x-axis depicts the stage at the time of nutrient addition. This strain displayed a statistically significant difference from wildtype cells in prometaphase I -anaphase I (*p<0.05, Fisher’s exact test) (G) Graph of the commitment assay for the *cdc5-as1* strain with CMK inhibitor (n = 108 cells; 3 independent experiments) and without inhibitor (n = 42 cells; 3 independent experiments). All cells were scored from nutrient addition at metaphase I (t = 0). Asterisks indicate statistically significant difference (**** p<0.0001, Fisher’s exact test). (H) Graph of the commitment assay for the *cdc5-as1 cdc20-mn* strain. CMK inhibitor added early (in pachytene; n = 140 cells; one experiment) and late (in metaphase I; n = 168 cells; 3 independent experiments). All cells were scored from nutrient addition in metaphase I. Asterisks indicate statistically significant difference (**** p<0.0001, Fisher’s exact test).

Because the *cdc5-mn* strains arrest with characteristics not typical of metaphase I, we asked if cells that undergo meiosis relatively normally, but that have reduced levels of Cdc5 could also have a defect in meiotic commitment. To this end, we analyzed commitment in a *CDC5/cdc5Δ* heterozygous strain. We found that 71% of *CDC5/cdc5Δ* cells were uncommitted when nutrients were added at metaphase I (Fig. 5F). In summary, our results demonstrate that Polo kinase has an essential role in meiotic commitment.

We considered that Cdc5’s role in meiotic commitment could occur during the normal meiotic process or could be triggered with nutrient addition. To distinguish between these possibilities, we studied strains with an inhibitable allele of *CDC5*, *cdc5-as1*(Attner, Miller, Ee, Elkin, & Amon, 2013; Snead et al., 2007). We first compared cells with and without the addition of the chloromethylketone (CMK) inhibitor. Like the *cdc5-mn* strains, addition of 5µM of CMK in prophase I caused a loss of commitment when nutrients were added at metaphase I (Fig. 5G). Without inhibitor, the cells remained committed to meiosis. Next, we asked whether Cdc5’s main role in meiotic commitment occurred prior to or at the time of nutrient-rich medium addition. We made *cdc5-as1* strains with a *cdc20-mn* to arrest cells in metaphase I. We compared cells with inhibitor added at prophase I (early) to cells that had inhibitor added after the metaphase I arrest but 30-minutes before nutrient addition (late). With inhibitor added late, cells are committed and remain in a metaphase I-like state, unlike the uncommitted cells with early addition of inhibitor (Fig. 5H). Because *CDC5* is expressed as cells exit prophase I, we conclude that Cdc5 establishes meiotic commitment in prometaphase I or metaphase I.

### Bmh1/Bmh2 Complex Activates Cdc5 to Ensure Meiotic Commitment

A previous study in human cells identified an interaction between Polo-like kinase 1 (Plk1) and the 14-3-3 isoform zeta that was important for Plk1’s role in cytokinesis (Du et al., 2012). This led us to question whether Cdc5 and Bmh1 can interact. Using a yeast two-hybrid assay, we identified an interaction between Bmh1 and Cdc5 (Fig. 6A).

**Figure 6:**
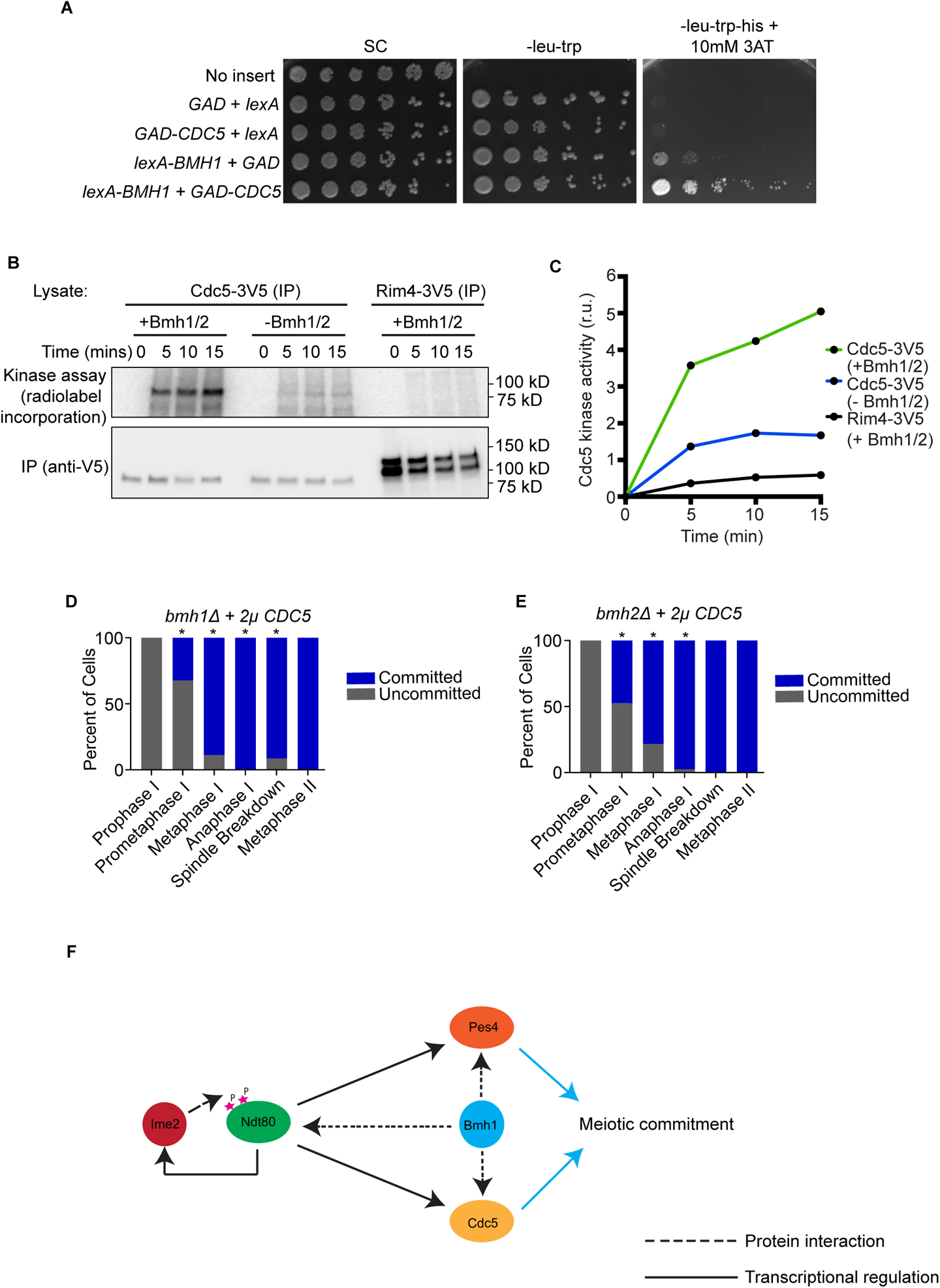
Bmh1 enhances Cdc5 kinase activity and regulate meiotic commitment. (A) Yeast-two hybrid strains were spotted onto the indicated plates with a 10-fold serial dilution. (B) Cdc5 kinase activity assay. Each assay contains immunoprecipitated of Cdc5-3V5 or Rim4- 3V5 as a control +/- purified Bmh1/2 complexes as indicated. Incorporation of radiolabeled ATP was assessed at the indicated times (upper). Immunoprecipitated Cdc5 and Rim4 are shown with the anti-V5 immunoblot (lower). (C) Graph depicting the radiolabeled ATP incorporation at the different timepoints. (D) Graph of the commitment assay for *bmh1Δ* + 2µ-*CDC5* strain. 222 cells were counted in two independent experiments. The x-axis shows the stage at the time of nutrient addition. This mutant displayed a statistically significant difference from *bmh1Δ* cells in prometaphase I - spindle breakdown (*p<0.05, Fisher’s exact test). (E) Graph of the commitment assay for the *bmh2Δ +* 2µ-*CDC5* strain. 175 cells were counted in two independent experiments. The x-axis shows the stage at the time of nutrient addition. This mutant displayed a statistically significant difference from *bmh2Δ* cells in prometaphase I, metaphase I, and anaphase I (*p<0.05, Fisher’s exact test). (F) Model for meiotic commitment. As cells exit prophase I, Ime2 phosphorylates Ndt80 and activates Ndt80. Ndt80 then transcribes *CDC5*, *PES4* and more *IME2*. Bmh1 interacts with Ndt80, Cdc5 and Pes4. Interaction of Bmh1 and Ndt80 enhances Ndt80 activity, interaction of Bmh1 and Cdc5 enhances Cd5 kinase activity, and Bmh1 interacts with Pes4 and establishes meiotic commitment.

Because numerous signaling pathways are regulated by 14-3-3 proteins, we considered that Bmh1 could modulate Cdc5 activity (van Hemert, Steensma, & van Heusden, 2001; van Heusden & Steensma, 2006). To test this hypothesis, we performed an *in vitro* kinase assay with immunoprecipitated Cdc5-3V5 with and without purified Bmh1/2 complexes. We chose to monitor autophosphorylation of Cdc5 as a measure of kinase activity (K. S. Lee & Erikson, 1997; Macurek et al., 2008; Mohapatra et al., 2015). We observed a low level of Cdc5 autophosphorylation in the absence of Bmh proteins (Fig. 6B,C). However, when Bmh1/Bmh2 complexes are added, Cdc5 autophosphorylation increases. Because we did not see a signal in our negative control (unrelated meiotic protein Rim4-V5), we can conclude that Cdc5 autophosphorylation we observed is not an artifact of the anti-V5 IP. These results suggest that Bmh1 enhances the activity of Cdc5.

We speculated that if Bmh1 also enhances Cdc5 activity *in vivo*, overexpression of *CDC5* could rescue meiotic commitment in *bmh1Δ* cells. We transformed *bmh1Δ* cells with a 2μ plasmid containing *CDC5*. Our commitment assays show an increased population of *bmh1Δ* committed cells when *CDC5* is overexpressed (Fig. 6D,2F; p<0.05, Fisher’s exact test).

Similarly, more *bmh2Δ* cells overexpressing *CDC5* display a committed phenotype compared to *bmh2Δ* without *CDC5* overexpression (Fig. 6E, 2G; p<0.05, Fisher’s exact test). These results suggest that increased activity of Cdc5 enhances meiotic commitment in *bmh1Δ* and *bmh2Δ* cells. We conclude that Bmh1/Bmh2 modulates Cdc5 activity for meiotic commitment. Overall, our results suggest a central role for 14-3-3 proteins in meiotic commitment.

## Discussion

Successful completion of meiosis requires that cells remain committed to completing meiosis, even in the presence of a mitosis-inducing signal. Historically, meiotic commitment has been studied in budding yeast due to the ease of providing meiotic cells a mitosis-inducing signal, nutrient-rich medium (Ballew & Lacefield, 2019a, 2019b; Esposito & Esposito, 1974; Gihana, Musser, Thompson, & Lacefield, 2018; Honigberg & Esposito, 1994; Nachman, Regev, & Ramanathan, 2007; Sherman & Roman, 1963; Simchen, 2009; Simchen et al., 1972; Tsuchiya et al., 2014). However, only a few genes have been implicated in meiotic commitment (Ballew & Lacefield, 2019a; Esposito & Esposito, 1974; Honigberg, Conicella, & Espositio, 1992; Honigberg & Esposito, 1994; Tsuchiya et al., 2014). Here, we provide a greater understanding of the meiotic commitment regulatory network by identifying five new components within the network: Bcy1, Ime2, Bmh1, Bmh2, and Cdc5. We show that these components are required to prevent cells from escaping meiosis and undergoing mitosis in the presence of nutrient-rich medium. Strikingly, our results suggest that commitment is actively maintained by several different proteins that function in multiple processes throughout meiosis. Our data allows us to propose a new model for how these components interact to maintain meiotic commitment (Fig. 6F). The 14-3-3 proteins are central to the model, in their role in regulating Ndt80 levels, Pes4, and Cdc5 activation.

Bcy1 represents a factor that ensures meiotic commitment by inhibiting the response to nutrient addition. Bcy1 is the regulatory subunit of PKA and restricts the activity of the catalytic subunits (Zaman, Lippman, Zhao, & Broach, 2008). In starved vegetative cells, the addition of glucose increases cAMP, which binds Bcy1, releasing the catalytic subunits for kinase activity (Thevelein & de Winde, 1999). PKA activity results in the expression of genes that increase ribosome production and glycolysis. A previous study describing the gene expression program of committed cells found that most sporulation-induced transcripts were downregulated upon nutrient-rich medium addition except *BCY1*, which was upregulated (Friedlander et al., 2006).

Furthermore, gene expression patterns that were altered in response to PKA activity in vegetative cells were not altered in committed cells, suggesting that committed cells are not responding to glucose addition. We showed that overexpression of *BCY1* increased the population of committed cells in the low Ndt80 strain (Fig. S1C). Conversely, loss of one copy of *BCY1* in an otherwise wildtype strain resulted in uncommitted cells (Fig. S1D). Therefore, our data combined with that of Friedlander *et al* suggest that the increased levels of Bcy1 in committed cells blocks the response to nutrient-rich medium and ensures meiotic commitment.

The other genes identified in the screen, *IME2, BMH1, BMH2,* and *CDC5* maintain meiotic commitment by ensuring the continuation of the meiotic program. Ime2, Bmh1, Bmh2, and Cdc5 have known roles in meiosis or sporulation, but only Ime2 had an obvious connection to meiotic commitment. Ime2 phosphorylates and activates Ndt80 to further induce transcription of the middle meiosis genes (Benjamin et al., 2003; Schindler et al., 2003; Sopko et al., 2002).

We showed that an *IME2* mutant allele with decreased activity had a strong commitment defect (Fig. S1G). Therefore, Ime2 is important for meiotic commitment likely through its role in activating Ndt80, which increases the levels of Ndt80 and Ndt80-dependent transcripts.

We identified Bmh1 and Bmh2 as key regulators of meiotic commitment, which interact with at least three other proteins important for meiotic commitment. First, we found that Bmh1 and Bmh2 modulate Ndt80 levels and Bmh1 and Ndt80 directly interact (Fig. 3A-G). Overall, these results suggested that the loss of meiotic commitment in *bmh1Δ* and *bmh2Δ* strains was due to low levels of Ndt80. In support of this conclusion, overexpression of NDT80 rescued meiotic commitment in *bmh1Δ* and *bmh2Δ* strains (Fig. 3H,I).

Second, we identified an interaction between Bmh1 and the putative RNA-binding protein Pes4. A prior study showed that Pes4 is needed for the delayed translation of approximately 30 Ndt80-dependent transcripts until meiosis II (Jin et al., 2017). Interestingly, unlike most Ndt80-induced mRNAs, a subset of the mRNAs bound by Pes4 were protected and the transcript levels did not drop with the addition of nutrient-rich medium (Friedlander et al., 2006; Jin et al., 2017). These RNAs lose protection in *pes4Δ* cells (Jin et al., 2017). Although *pes4Δ* cells underwent meiosis and sporulation normally, we showed that a large population of *pes4Δ* cells failed to maintain meiotic commitment (Fig. 4A). In addition, overexpression of *PES4* increases the population of committed *bmh1Δ* cells (Fig. 4B). Combined, these results suggest that Pes4 maintains meiotic commitment, possibly by protecting transcripts whose encoded protein products are used in meiotic exit and sporulation.

Finally, we found that Bmh1 enhances the autophosphorylation of Cdc5, which further activates Cdc5 (Fig. 6B,C). Loss of Cdc5 activity, either with a meiotic null allele or with an inhibitable allele, causes a loss of meiotic commitment (Fig. 5D-G). An interesting future direction is to identify the substrates of Cdc5 needed for meiotic commitment. We show that the Cdc5 activity that protects meiotic commitment is needed during prometaphase/metaphase I, prior to nutrient-rich medium addition (Fig. 5H). These results suggest that a normal meiotic role of Cdc5 is also essential for meiotic commitment.

Our results reveal a model for a meiotic commitment regulatory network in which Bmh1 and Bmh2 are central regulators that prevent cells from escaping meiosis in the presence of nutrient-rich medium (Fig. 6F). As cells exit prophase I, Ime2 activates Ndt80, which leads to the transcription of *CDC5*, *PES4,* and more *IME2*. We show that Bmh1/Bmh2 interact with three of these proteins required for meiotic commitment: Ndt80, Pes4, and Cdc5. The interaction between Bmh1/2 and Cdc5 leads to enhanced kinase activity. We hypothesize that the interaction between Bmh1 and Pes4 leads to the proper folding or stability of Pes4 to protect mRNAs from down-regulation and ensure their translation at the end of meiosis, allowing normal meiotic exit and spore formation. Intriguingly, this regulatory network only becomes essential for meiotic maintenance in the presence of nutrient-rich medium. Therefore, this work raises the exciting possibility that the temporal regulation of translation by an RNA-binding protein has evolved to ensure cells commit to completing meiosis when exposed to a mitosis-inducing signal.

## Materials and Methods

### Strains

Unless noted, *Saccharomyces cerevisiae* strains used in this study are derivatives of W303 (*ade2-1 can1-100 leu2-3, 112 his3-11, 15 ura3-1 trp1-1*) and can be found in Supplementary Table S3. The strains used for the yeast two-hybrid experiment were derivatives of L40 (*his3Δ200 trp1-90 leu2-3,112 ade2 lys2::lexA_op_-HIS3::LYS2 gal80 ura3:: lexA_op_- lacZ::URA3*) with the *GAD* and *lexA* plasmids transformed in. Standard PCR-based methods were used for deleting and tagging genes and swapping promoters (Janke et al., 2004; Longtine et al., 1998). The low Ndt80 strain, with the deletion of the MSEs in the promoter, was from a previous study (Tsuchiya et al., 2014). The *mec1Δ sml1Δ* mutations were introduced in the strain LY6981 by crossing in and mating subsequent haploids. The *CDC5/cdc5* heterozygous strain was created by deleting *CDC5* in a diploid strain. The haploid *cdc5-as1* strain carrying *cdc5L158G* was a gift from the Marston lab and crossed into strains. The *bmh1-3A* strain was constructed by integrating pLB519 in the haploid *bmh1Δ* strains at the *BMH1* promoter.

### Plasmids

Oligos and plasmids can be found in Tables S4 and S5, respectively. Empty 2µ vector (pLB227) was made from the A4 plasmid from the Yeast Genomic Tiling Collection by digesting with SpeI and NheI and religating the 6.5kb fragment (G. M. Jones et al., 2008). *P_NDT80_-NDT80:LEU2* (pLB225) was made by digesting the A6 plasmid from the Yeast Genomic Tiling Collection using SalI and SacI and the 7.1 kb band was ligated into cut pLB107. *P_BMH1_- BMH1:LEU2* (pLB262) was made by amplifying *P_BMH1_-BMH1* using oligos LO1677 and LO1678. The fragment was digested with SalI and BamHI and ligated with cut YEplac181. *P_BMH2_-BMH2:LEU2* (pLB539) was made by amplifying *P_BMH2_-BMH2* using oligos LO1968 and LO1969, digesting with SalI and XmaI and ligating into cut YEplac181. *lexA-BMH1:TRP1* (pLB518) was made by amplifying *BMH1* ORF using oligos LO3089 and LO3090, digesting with EcoRI and SalI and ligating into cut pBTM116. *GAL4AD-CDC5:LEU2* (pLB491) was made by amplifying the *CDC5* ORF using LO3062 and LO3063 and ligating with NcoI and XhoI digested pACTII. *P_CDC5_-CDC5:URA3* (pLB465) and *P_CDC5_-CDC5:LEU2* (pLB463) were made by amplifying the *P_CDC5_-CDC5* ORF with oligos LO2800 and LO2801, and ligating into BamHI and SacI digested YEPlac195 and YEplac181 respectively. *P_BMH1_-BMH1* was amplified with LO1677 and LO1678, and cloned in SalI and BamHI digested pRS403 (ref Sikorski and Heiter, 1989). *P_BMH1_-bmh1-3A:HIS3* (pLB519) was made by Genewiz, using *P_BMH1_-BMH1:HIS3* (pLB509) as a template. *P_PES4_-PES4:URA3* (pLB513) was made by amplifying *P_PES4_-PES4* with oligos LO3163 and LO3164 and ligating in SalI and SacI digested YEplac195. *P_BCY1_- BCY1:LEU2* (pLB306) was made by amplifying *P_BCY1_-BCY1* with oligos LO1808 and LO1809 and ligating in SalI and BamHI digested YEplac181. *P_IME2_-IME2:LEU2* (pLB258) was made by amplifying *P_IME2_-IME2* with oligos LO1673 and LO1674 and ligating into SalI and XmaI digested YEplac181. *P_IME2_-IME2-myc-TRP1* (pLB268) was made by amplifying *P_IME2_-IME2- myc* using oligos LO686 and LO687. *P_IME2_-IME2 T242A-myc-TRP1* (pLB285) was made by using pLB268 as a template and oligos LO1627 and LO1629 for site-directed mutagenesis.

### Media

The following yeast media were used in this study: YPD (2% peptone, 1% yeast extract, 2% glucose), YPA (2% peptone, 1% yeast extract, 2% potassium acetate), SPM (1% potassium acetate), SCD –leu (0.67% YNB without amino acids, 0.2% dropout mix containing all amino acids except leucine, 2% glucose), SCA –leu (0.67% YNB without amino acids, 0.2% dropout mix containing all amino acids except leucine, 2% potassium acetate), SCD –ura (0.67% YNB without amino acids, 0.2% dropout mix containing all amino acids except uracil, 2% glucose), SCA –ura (0.67% YNB without amino acids, 0.2% dropout mix containing all amino acids except uracil, 2% potassium acetate), SCD-leu-trp (0.67% YNB without amino acids, 0.2% dropout mix containing all amino acids except leucine and tryptophan, 2% glucose), and 2XSC (1.34% YNB without amino acids, 0.4% dropout mix containing all amino acids, 4% glucose).

### Commitment assay

Cells were grown in YPD for 16-18 hours at 30°C, transferred to YPA (1:50 dilution) for 12-14 hours at 30°C, washed once with sterile water, resuspended in SPM and incubated on a roller drum at 25°C for 9:30 hours (except the low Ndt80 strain background was incubated for 13:30 hours and *cdc20mn*, *cdc5mn* and *cdc5-as1* strains were incubated for 12-14 hours). Cells were then loaded into the microfluidics chambers (CellAsic Y04D yeast perfusion plates). For inhibition of Cdc5 in the *cdc5-as1* strain, 5µM chloromethylketone (CMK) inhibitor (gift of A. Marston) was added in the culture tube 10 minutes prior to loading and was also added to 2XSC. SPM flowed through the chamber for 20 minutes and cells were exposed to 2XSC 10hrs after SPM addition, (14hrs for the low Ndt80 strain background and 12-14 hrs for the *cdc20mn*, *cdc5mn* and *cdc5-as1* strains). The commitment assay was performed using the CellAsic Onix microfluidics perfusion platform.

### Microscope image acquisition and time-lapse microscopy

Cells were imaged using a Nikon Ti-E inverted microscope equipped with SNAPHQ2 CCD camera (Photometrics), 60X oil objective (PlanAPO VC, 1.4NA). For each experiment, 30 random fields were selected. Five z-steps (1.2µm) were acquired for each field. Images were acquired in 10 minutes interval for 8-12 hours. The exposure times for Brightfield was 60ms, 500-700ms for GFP and 700-900ms for mCherry. Z-stacks were combined into a single maximum intensity projection with NIS-Elements software (Nikon).

### Overexpression screen

The overexpression screen was performed in 96-well dishes. The low Ndt80 strain was transformed with the Yeast Genomic Tiling Collection (G. M. Jones et al., 2008) in 96-well dishes using the following protocol (Gietz, 2014): Cells were grown in 20mL of YPD for 12 hours at 30°C. 2.5 x 10^8^ cells were transferred to 50ml of pre-warmed YPD and incubated at 30°C for 4 hours. 200uL of the cell suspension was transferred to 96-well plates and the plates were centrifuged for 10 minutes at 1300g. The supernatant was discarded. 5µL of the plasmids from the Yeast Genomic Tiling Collection was placed in each well. 35µL of the transformation mix (15µL of 1M Lithium acetate + 20µL of boiled 2mg/mL salmon sperm DNA) and 100µL of 50% PEG (MW 3350) was added to the cell pellet and incubated at 42°C for 2 hours. The plate was centrifuged at 1300g for 10 minutes, 10µL of sterile water was added to the cells and 5µL of cells were spotted on SC-leu plates. The plates were incubated at 30°C for 2-4 days.

For commitment assays in 96-well dishes, 1mL of SCD was added to each well in a sterile 96-well block. A single colony was inoculated and incubated in the 96-well blocks at 30°C for 24 hours. 40uLs was then diluted into 1mL of SCA. The wells contained sterile magnetic stirrers (3mm x 5mm) and the block was incubated at 30°C for 15hrs on a shaker. The block was covered with sterile air permeable covers. The block was spun at 3000RPM for 1 minute. The supernatant was removed, the cells were washed twice with 1mL of sterile water and then resuspended in 1mL of SPM. The block was sealed with an air permeable seal and stirred at 25°C for 20 hours. The block was spun again at 3000RPM for 1 minute and the supernatant was removed and cells were resuspended in 1mL of 2XSC. The block was placed on the magnetic stirrer at 25°C for 8hrs. The block was centrifuged at 3000RPM for 1 minute. The supernatant was discarded and the cells were transferred to a 96-well round bottom plate. The cells were fixed in 150uL of freshly prepared 4% para-formaldehyde.

For imaging in 96-wells, 5µL of the fixed cells were added to each well. To get a monolayer of cells, an agar pad made up of 1.2% agarose in 1XPBS (pH7.4) was placed on the cells. Cells were imaged at room temperature using a Nikon Eclipse Ti inverted microscope equipped with Hammamatsu Orca-Flash4.0 sCMOS camera 60X oil objective (PlanAPO VC, 1.2NA). For each well, 10 random fields were selected. Five z-steps (1.2µm) were acquired for each well. GFP-Tub1 and Spc42-mCherry were imaged with a 500ms exposure and 1 ND filter. Brightfield images were acquired with a 60ms exposure.

### Western blotting

For Western blotting analysis of Ndt80, Cdc5, Clb1 and Pgk1, protein extraction was carried out by using a TCA method (Wang et al., 2020). 5mLs of meiosis culture was harvested for each timepoint. Cells were spun at 3000RPM for 1 minute. The supernatant was removed and 5mL of ice-cold 10% Trichloroacetic acid (TCA) was added for 10 minutes. Cells were centrifuged at 5000RPM for 1 minute. The supernatant was removed and 1mL of ice-cold acetone was added to the cells. Cells were vortexed and centrifuged at 14000RPM for 1 minute. The supernatant was discarded and this step was repeated twice. Tubes were left open to dry in the hood for 3 hours. For the protein extraction, 200µL of protein breakage buffer (60mM Tris pH 7.5, 1.2mM EDTA pH8, 3.3 mM DTT, Pierce Protease Inhibitor Mini tablet-EDTA free) was added along with 200µL of 0.5mm glass beads. Cells were broken by vortexing six times with 1- minute pulses, with 1 minute on ice in between. 100µL of 3X SDS buffer was added and tubes were boiled for 5 minutes. The supernatant was resolved on SDS PAGE gel at 150V for 90 minutes and transferred onto a PVDF membrane. The membranes were blocked with 5% milk in 1X TBST.

The primary antibodies used were a rabbit anti-Ndt80 (generous gift of M. Lichten, 1:10000), anti-Pgk1 mouse (Invitrogen, Catalog #459250, 1:10000), anti-Cdc5 goat (Santa Cruz Biotechnology, sc-6733, 1:1000), anti-Clb1 goat (Santa Cruz Biotechnology, sc-7647, 1:1000). The secondary antibodies used were ECL-anti-rabbit HRP IgG (GE Healthcare, NA9340V, 1:10000), ECL-anti-mouse HRP IgG (GE Healthcare, NA9310V, 1:10000), anti-goat IgG HRP (R&D Systems, HAF109, 1:5000).

Densitometry analysis of each protein was carried out by using ImageJ software. The individual densitometry values of Ndt80, Cdc5 and Clb1 were divided by the densitometry values of Pgk1 for the Ndt80/Pgk1, Cdc5/Pgk1 and Clb1/Pgk1 ratios. The graphs were generated by using GraphPad Prism software.

### Yeast-two hybrid assay

Transformants containing the *GAD* plasmids and *lexA* plasmids were grown in SCD-leu-trp at 30°C for 16-18 hours. LY8522 was grown in SC for 16-18 hours. The cells were diluted 1:10 in sterile water and the optical density at 600nm (OD_600_) was measured using a spectrophotometer. Culture volumes equivalent to 0.1 OD were set as starting dilution and were further subjected to 10-fold serial dilution. The serial dilutions were then spotted on SC plates, SC –leu-trp plates and SC –leu-trp-his + 10mM 3-aminotriazole (Sigma) plates. The plates were incubated at 30°C for 3-5 days.

### 14-3-3 C-terminus sequence analysis

The following FASTA sequences were obtained from NCBI protein resource: NP_011104.3, NP_010384.3, NP_594167.1, NP_564249.1, NP_565977.1, NP_001014516.1, NP_001129171.1, NP_724887.2, NP_001240734.1, NP_003395.1, XP_005158144.1, NP_001076267.1 and NP_061223.2. The sequences were aligned using blastp suite. The sequences around *Saccharomyces cerevisiae* Bmh1 S235, S238 and S240 were aligned with the 14-3-3 protein sequences across different organisms.

### Immunoprecipitation and mass spectrometry analysis

Cells were grown for 16-18 hours in 2mL YPD, diluted 1:50 dilutions into 50mL YPA, and incubated at 30°C for 12-14 hours. Cells were washed once with sterile water and resuspended in 50mL SPM and incubated at 25°C. After 10hrs in SPM, cells were spun down and snap frozen in liquid nitrogen. The cell pellet was thawed on ice, 200µL NP40 lysis buffer (50mM Tris pH7.5, 150mM NaCl, 2mM MgCl2, 1%NP-40, 10% glycerol, Pierce Protease Inhibitor Mini tablet-EDTA free) and 200µL of 0.5mm glass beads were added. The cells were lysed by bead beating 6 times for 1min with 1min on ice in between. The tubes were centrifuged at 4°C at 18000 RCF for 10mins. The supernatant was removed and the protein concentration was measured by Bradford assay (Pierce Coomassie Plus Assay kit, Thermo Scientific, Catalog #23236). 25µL of DYNAbeads Protein G (Invitrogen, Catalog # 10003D) was equilibrated for each IP. 1mg of protein, 10µg anti-GFP mouse (Roche, Catalog #11814460001) and 1mL of NP-40 lysis buffer was added to the DYNAbeads tube. IP samples were incubated on a rotor at 4°C for 12 hours. The beads were immobilized by placing the tube against a magnetic bar. The supernatant was removed and the beads were washed with 1mL of NP-40 lysis buffer. The wash steps were repeated twice and 25µL of 3X SDS buffer was added to the beads. The beads were boiled for 5mins. The supernatant was loaded in a 10-well 4-20% Mini-PROTEAN TGX Precast protein gel (BIO-RAD, Catalog #4561094). The gel was run at 150V for 5 minutes. After the mini-run, the gel was washed 3 times for 10 minutes with distilled water. The gel was stained with Imperial Protein Stain (Thermo Fisher, Catalog #24615) for 30 minutes. To remove excess stain, the gel was washed 3 times for 10 minutes in distilled water. The protein band was excised from the gel and analyzed by mass spectrometry.

Gel bands were diced with a razor blade. Samples were washed twice in 50% acetonitrile, 0.1% formic acid and dried down in a SpeedVac concentrator (Thermo Scientific). Disulfide bonds were reduced by incubation for 45 min at 57 °C with a final concentration of 10 mM Tris (2-carboxyethyl) phosphine hydrochloride (Catalog no C4706, Sigma Aldrich). The gels were washed and a final concentration of 20 mM iodoacetamide (Catalog no I6125, Sigma Aldrich) was then added to alkylate these side chains and the reaction was allowed to proceed for one hour in the dark at 21 °C. The gels were washed and dried down. Gel pieces were rehydrated in 25 mM ammonium bicarbonate containing 200 ng of trypsin (V5113, Promega) and the samples were digested for 14 hours at 37 °C. The following day, peptides were extracted twice using 50% acetonitrile in 0.1% formic acid. The extracts were dried down and desalted using zip tips (EMD Millipore).

Samples were analyzed by LC-MS on an Orbitrap Fusion Lumos (ThermoFisher) equipped with an Easy NanoLC1200 HPLC (ThermoFisher). Peptides were separated on a 75 μm × 15 cm Acclaim PepMap100 separating column (Thermo Scientific) downstream of a 2 cm guard column (Thermo Scientific). Buffer A was 0.1% formic acid in water. Buffer B was 0.1% formic acid in 80% acetonitrile. Peptides were separated on a 60 minute gradient from 0% B to 35% B. Peptides were collisionally fragmented using HCD mode. Precursor ions were measured in the Orbitrap with a resolution of 120,000. Fragment ions were measured in the ion trap. The spray voltage was set at 2.2 kV. Orbitrap MS1 spectra (AGC 1×106) were acquired from 400- 1800 m/z followed by data-dependent HCD MS/MS (collision energy 42%, isolation window of 0.7 Da) for a three second cycle time. Charge state screening was enabled to reject unassigned and singly charged ions. A dynamic exclusion time of 60 seconds was used to discriminate against previously selected ions.

The LC-MS/MS data was searched using Proteome Discoverer 2.1. MS spectra were searched against the *Saccharomyces cerevisiae* database downloaded from Uniprot on 4/2018. The database search parameters were set as follows: two missed protease cleavage sites were allowed for trypsin digested with 10 ppm precursor mass tolerance and 0.6 Da for fragment ion quantification tolerance. Oxidation of methionine, pyroglutamine on peptide amino termini and protein N-terminal acetylation were set as variable modifications. Carbamidomethylation (C; +57Da) was set as a static modification.

### Cdc5 immunoprecipitation/kinase assay

25 ml cultures were pelleted, washed once with Tris (pH 7.5), transferred into a 2 ml tube and snap frozen in liquid nitrogen for later processing. Cells were broken with Zirconia/Silica beads in 200 µL NP40 Lysis Buffer (50 mM Tris pH 7.5, 150 mM NaCl, 1% NP-40, 10% glycerol) containing 1mM DTT and 1x Halt protease and phosphatase inhibitors (Thermo). After breaking, extracts were cleared twice by centrifugation and protein concentration was determined by Bradford assay. Immunoprecipitations were performed in 1 ml diluted extract (800 µg total protein). Cdc5-3V5 and Rim4-3V5 (control non-kinase tagged protein) were immunoprecipitated at 4°C 2 hours using 20 µL of anti-V5-agarose slurry (Sigma). Purifications were washed 4 times with NP40 buffer.

For the kinase reaction, beads were incubated in kinase buffer (25 mM Tris-HCl pH 7.5, 5 mM beta-glycerophosphate, 1 mM dithiothreitol (DTT), 0.1 mM Na_3_VO_4_, 10 mM MgCl_2_, 10 mM ATP (non-radioactive) alone or with 8 µM Bmh1/Bmh2 protein complexes purified from yeast. The reaction was initiated by adding 1 ul of [γ-^32^P] ATP (3000Ci/mmol) to each sample. 5 ul samples were withdrawn after the indicated amount of time. To stop the reaction, 2.5 ul 3 x SDS loading Buffer (9% SDS, 0.75 mM Bromophenol blue, 187.5 mM Tris-HCl pH 6.8, 30% glycerol, and 810 mM β-mercaptoethanol) was added and samples were boiled for 5 mins.

Samples were separated on a 10% SDS-PAGE gel and transferred to a nitrocellulose membrane using a semi-dry transfer (Biorad). The membrane was dried, mounted on a phosphor screen, and imaged on a Typhoon imager (GE Healthcare). Immunoprecipitations were analyzed by detection with anti-V5.

### Statistical analysis

All statistical analyses were performed using GraphPad Prism software. For all microscopy data, the individual cell numbers were entered in the contingency tables and two-sided Fisher’s exact test was used to determine the significance. The total number of cells analyzed in each graph are indicated in the figure legends. For the protein quantification in figure 3, statistical significance was determined for t = 14 using an Unpaired t-test with Welch’s correction. For the protein quantification in Supplementary Figure 3, statistical significance was determined using Two-way Anova. Differences among compared data was considered statistically significant if the p-value was <0.05.

## Acknowledgements

We thank Nancy Hollingsworth, Angelika Amon, Michael Lichten, and Adele Marston for strains and reagents. We thank Rolf Sternglanz for discussions. We thank the Lacefield lab for insightful comments on the manuscript. We thank the Light Microscopy and Imaging Center at Indiana University, especially Jim Powers for assistance. This work was supported by a grant from the NIH (GM105755).

## Supplementary Figure Legends

**Supplementary Figure 1:**
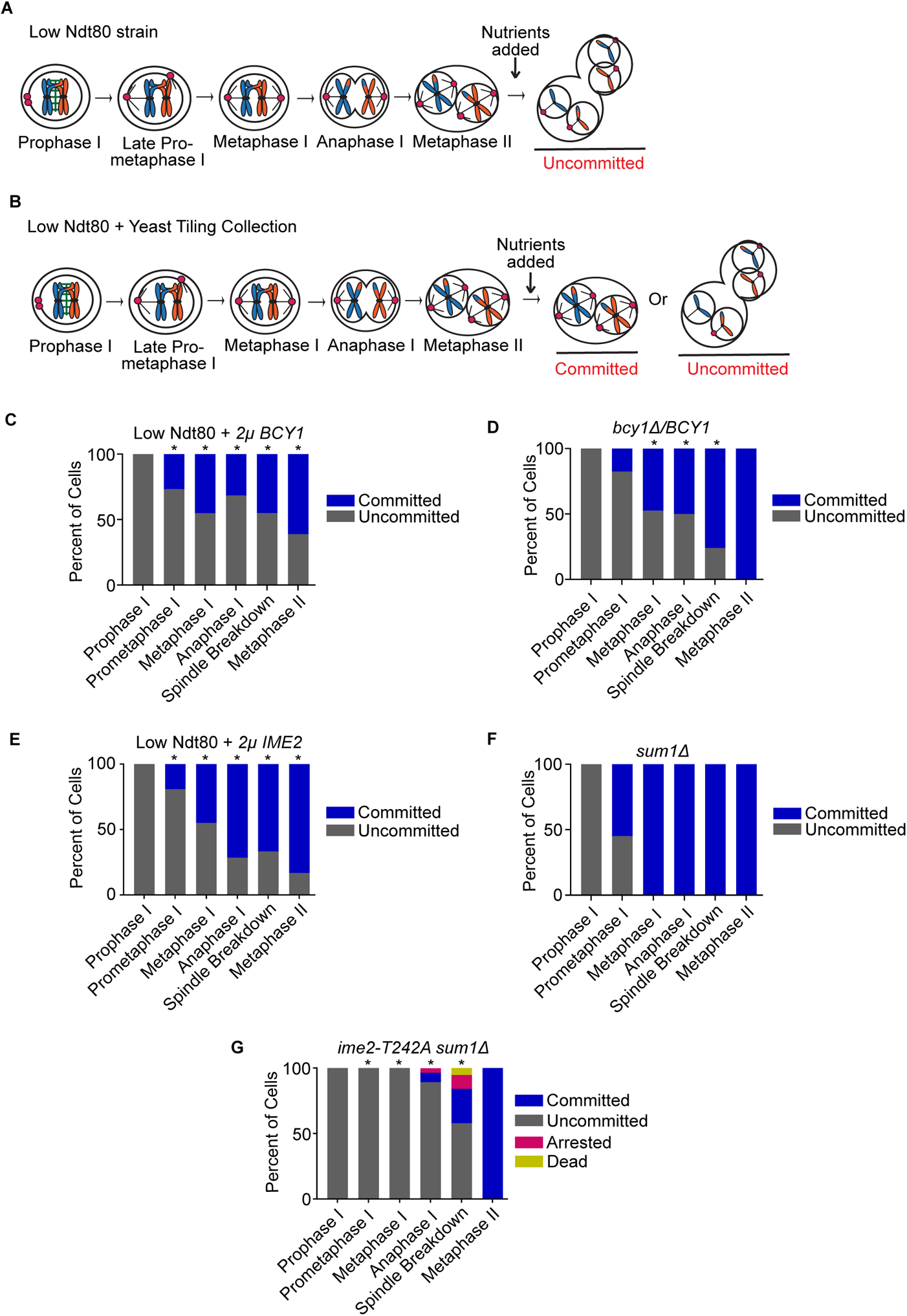
*BCY1* and *IME2* are important for meiotic commitment. (A,B) Cartoon for the overexpression screen. (C) Graph of the commitment assay for the low Ndt80 strain with 2µ*-BCY1*. 194 cells in two independent experiments were counted. The x-axis depicts the stage at the time of nutrient addition. This mutant displayed a statistically significant difference from Low Ndt80 cells in prometaphase I - metaphase II (*p<0.05, Fisher’s exact test). (D) Graph of the commitment assay for *bcy1Δ*/*BCY1* heterozygous cells. 108 cells were counted over two independent experiments. The x-axis depicts the stage at the time of nutrient addition. This mutant displayed a statistically significant difference from wildtype cells in metaphase I, anaphase I and spindle breakdown stages (*p<0.05, Fisher’s exact test). (E) Graph of the commitment assay for the low Ndt80 + 2µ-*IME2* strain. 231 cells in two independent experiments were counted. The x-axis depicts the stage at the time of nutrient addition. This mutant displayed a statistically significant difference from the low Ndt80 strain in prometaphase I - metaphase II stages (*p<0.05, Fisher’s exact test). (F) Graph of the commitment assay for the *sum1Δ* strain. 125 cells were counted in two independent experiments. The x-axis depicts the stage at the time of nutrient addition. This mutant is not significantly different from wildtype cells (Fisher’s exact test). (G) Graph of the commitment assay for the *ime2-T242A sum1Δ* cells. 147 cells were counted over four independent experiments. The x-axis depicts the stage at the time of nutrient addition. This mutant displayed a statistically significant difference from wildtype cells in prometaphase I - spindle breakdown stages (*p<0.05, Fisher’s exact test).

**Supplementary Figure 2:**
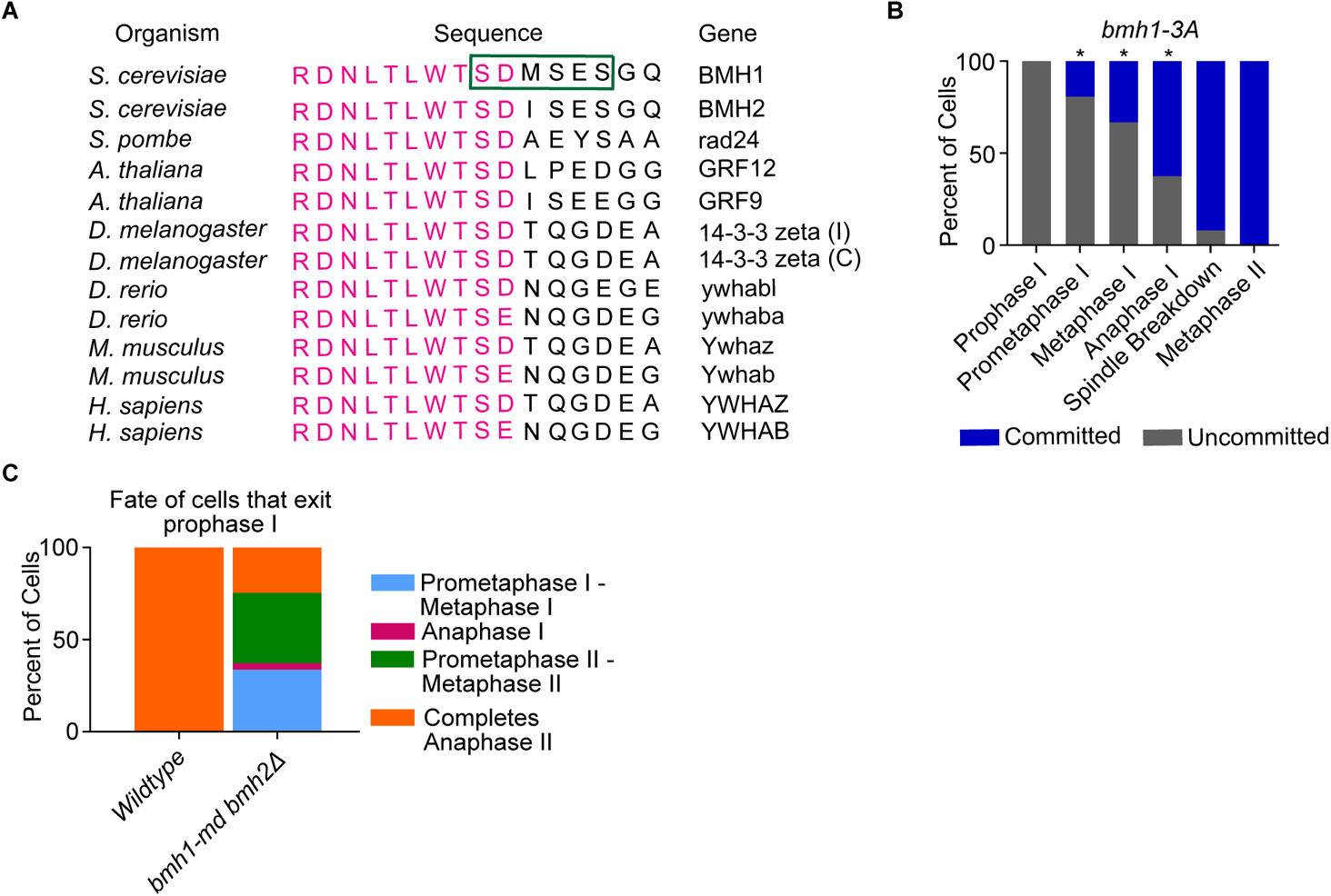
Cells with a mutation in a highly conserved residue in Bmh1 have a commitment defect and cells lacking both Bmh1 and Bmh2 fail to complete meiosis. (A) The protein sequence around the S235, S238 and S240 residues of Bmh1 were aligned with the 14-3-3 protein sequences across different organisms, as indicated. Magenta denotes conserved sequences. The green box highlights amino acids 235 to 240 in Bmh1. (B) Graph of the commitment assay for the *bmh1-3A* cells. 162 cells were counted over two independent experiments. The x-axis depicts the stage at the time of nutrient addition. This mutant displayed a statistically significant difference from the wildtype strain in prometaphase I, metaphase I, and anaphase I (*p<0.05, Fisher’s exact test). (C) Graph depicting the fate of cells that exit prophase I in the *bmh1-md bmh2Δ* strain. This graph shows the 99% of the wildtype cells and 20% of *bmh1-md bmh2Δ* cells that exit prophase I.

**Supplementary Figure 3:**
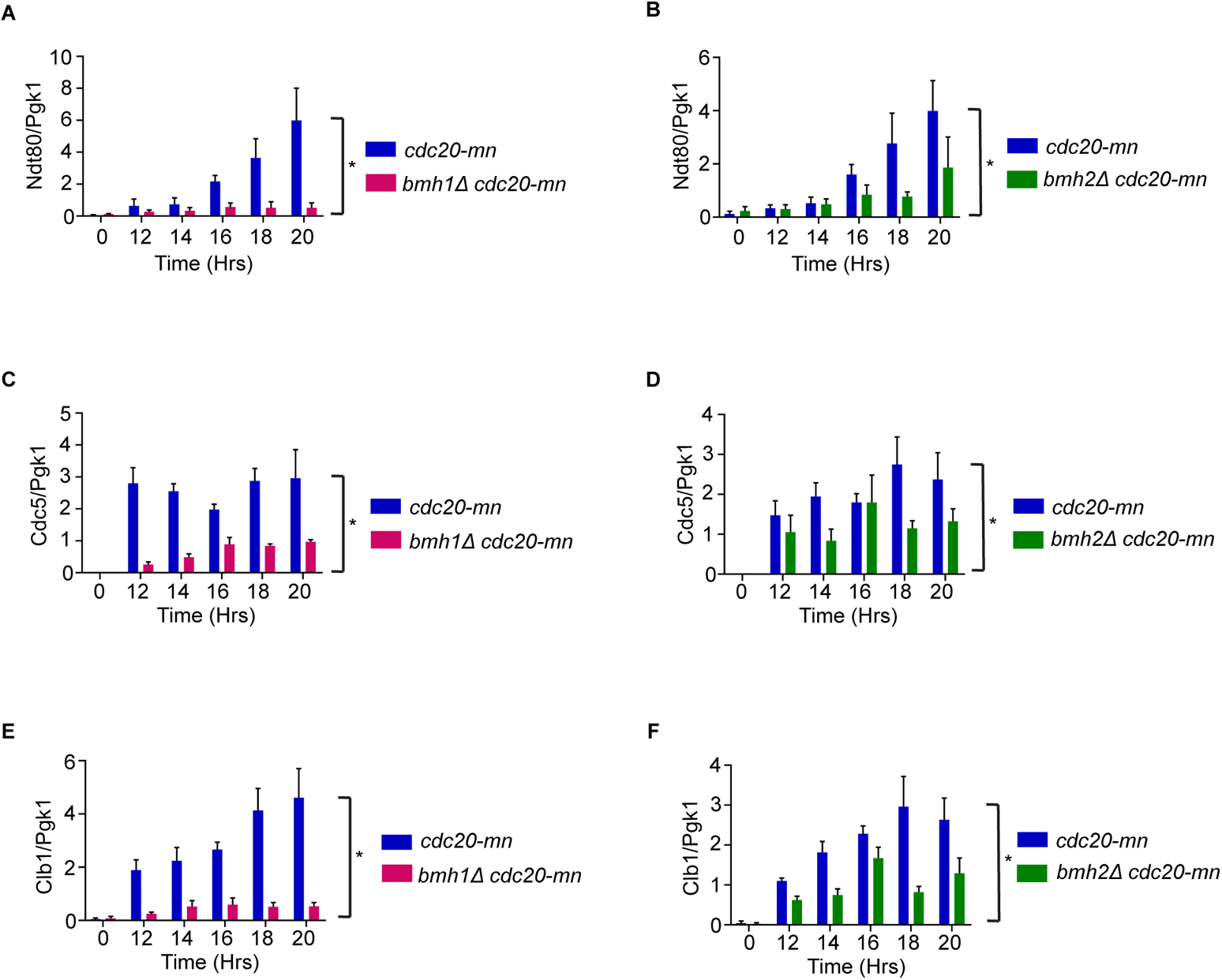
Quantification of the Ndt80, Cdc5, and Clb1 levels in *cdc20mn*, *bmh1Δ cdc20mn*, and *bmh2Δ cdc20mn* cells. Quantification of Ndt80 levels (A and B), Cdc5 levels (C and D), Clb1 levels (E and F) in *cdc20mn*, *bmh1Δ cdc20mn,* and *bmh2Δ cdc20mn* cells. Time zero indicates the time at which cells were transferred to SPM medium. Graphs shows mean ± SEM of the densitometric analysis of Ndt80 levels/Cdc5 levels/Clb1 levels relative to Pgk1 levels at the indicated time points. Three independent experiments were performed for each genotype. Asterisks show statistical significance (* p<0.05, Two-way Anova).

