## Supplemental Table S1-S5 for "The 14-3-3 Proteins Bmh1 and Bmh2 Are Key Regulators of Meiotic Commitment"

**Supplemental Table S1.** Plasmids from the Yeast-tiling collection that rescued commitment defect in Low Ndt80 strain

| Clone | Genes |
|---|---|
| YGPM27k09 | *[DEP1]&, CYS3, SWC3, MDM10, SPO7, FUN14, ERP2, tP(UGG)A, [SSA1]&* |
| YGPM7c11 | *[ROX3]*, RPL32, SCS22, MAP2, MRP21, AVT5, [TEL1]&* |
| YGPM24k20 | *[HHF1]*, HHT1, IPP1, YBR012C, YBR012W-A, [YBR012W-B]** |
| YGPM26d06 | *SCO1, CHS2, ATP3, FIG1, FAT1, [YBR042C]&* |
| YGPM2o04 | *[YBR063C]*, [YBR064W], ECM2, NRG2, TIP1, BAP2, TAT1, YBR070C, YBR071W, HSP26, YBR072C-A, RDH54, [YBR074W]** |
| YGPM12l03 | *[MMS4]&, YBR099C, FES1, EXO84, SIF2, YBR103C-A, YMC2, [VID24]&* |
| YGPM22d16 | *[HIS7]*, ARO4, YBR250W, MRPS5, DUT1, SRB6, TRS20, [YBR255W]** |
| YGPM22o22 | *[YCL019W]&, tL(CAA)C, LEU2, NFS1, DCC1, [BUD3]** |
| YGPM28o02 | *[BPH1]&, SNT1, FEN1, RRP43, RBK1, [PHO87]&* |
| YGPM23o21 | *[CPR4]&, IMG2, YCR072C, SSK22, SOL2, ERS1, YCR075W-A, [YCR076C]&* |
| YGPM32d08 | *[SSB1], YDL228C, HO, GCS1, SHS1, [WHI4]&* |
| YGPM14d22 | *GCS1, SHS1, WHI4, HBT1, FMP45, YDL221W, [CDC13]&* |
| YGPM14m05 | *[YDL221W]&, [CDC13], DTD1, YDL218W, TIM22, RRI1, [GDH2]&* |
| YGPM20d10 | *OSH2, ERP3, CDC7, YDL016C, TSC13, NOP1, [HEX3]** |
| YGPM23m11 | *[GRX3]*, tQ(UUG)D2, YDR098C-B, YDR098C-A, BMH2, TVP15, ARX1, YDR102C, STE5, [SPO71]&* |
| YGPM2l13 | *[SEC1]*, TRM82, SEC5, TAF10, CDC37, STB3, YDR169C-A* |
| YGPM12l05 | *LPP1, SPG3, PSP1, YDR506C, tL(CAA)D, [GIN4]&* |
| YGPM26m21 | *[TRP2]&, [YER090C-A], MET6, YER091C-A, IES5, TSC11, [YER093C-A]* |
| YGPM12l10 | *[RAD24]&, GRX4, TMT1, YER175W-A, ECM32, BMH1, [PDA1]* |
| YGPM26n21 | *[POP6]*, [YGR031W], YGR031C-A, GSC2, FMP17, RPL26B, YGR035C, YGR035W-A, [CAX4]* |
| YGPM14o13 | *[CLB1], CLB6, tC(GCA)G, YGR109W-A, YGR109W-B* |
| YGPM20c13 | *[NDT80], tF(GAA)H2, YHR125W, YHR126C, YHR127W, FUR1, ARP1,* |

| | |
|---|---|
| | *YHR130C, [YHR131C]&* |
| YGPM5m24 | *[VTH1]&, YIL172C, YIL171W-A, YIL171W, YIL170W* |
| YGPM7n05 | *[CKA1]*, CAP2, BCY1, YIL032C, ULP2, YIL030W-A, [SSM4]&* |
| YGPM20a02 | *[UTP10]*, PRM10, YJL107C, IME2, SET4* |
| YGPM16m18 | *[JEM1]&, PSF2, ARG2, YJL070C, UTP18, YJL068C, [YJL067W], [MPM1]&* |
| YGPM13p14 | *[YJL067W], MPM1, DLS1, YJL064W, MRPL8, YJL062W-A, LAS21, NUP82, [BNA3]** |
| YGPM3e15 | *[PRP16]&, OMA1, TVP38, STC1, PXL1, SRL3, SRP40, [PTR2]** |
| YGPM8g21 | *[UBI4]*, ENT4, YLL037W, PRP19, GRC3, RIX7, YLL033W, [YLL032C]&* |
| YGPM7a09 | *[IMD4]*, [snR54]*, SPC2, CYB2, YML054C-A, tY(GUA)M1, YML053C, SUR7, GAL80, YML050W, [RSE1]&* |
| YGPM5d16 | *[YML002W]&, YPT7, CDC5, YMR001C-A, YMR002W, YMR003W, [MVP1]** |
| YGPM30k17 | *[YMR110C]*, YMR111C, MED11, FOL3, YMR114C, FMP24, ASC1, snR24, SPC24, [YMR118C]* |
| YGPM21j12 | *[PEP5], FUS2, YMR233W, RNH1, RNA1, TAF9, [BCH1]** |
| YGPM19l23 | *[YNL300W], TRF5, CLA4, [MON2], YNL296W* |
| YGPM10o14 | *[SIP3]*, [FOL1], GIS2, YNL254C, TEX1, [MRPL17]* |
| YGPM12g06 | *[YNL228W], JJJ1, YNL226W, CNM67, YNL224C, ATG4, SSU72, [snR19], [POP1]&* |
| YGPM10e13 | *[SIS1]*, LST8, MRP7, HRB1, PET8, RLP7, DOM34, CIT1, YNR001W-A, tP(AGG)N, tN(GUU)N2, [FUN34]* |
| YGPM2k08 | *[PPG1]&, HUB1, ABZ1, SOL1, YNR034W-A, ARC35, YNR036C, RSM19, DBP6, [ZRG17]&* |
| YGPM15l07 | *[YOL097W-A]&, WRS1, COQ3, HMI1, RFC4, TRM10, YOL092W, SPO21, MSH2, [HAL9]&* |
| YGPM32g14 | *[IRA2]&, REX4, YOL079W, [AVO1]* |
| YGPM3b12 | SLY41, SNU66, YOR309C, NOP58, HSD1, RPL20B, SPS4, YOR314W, YOR314W-A, YOR315W, [COT1]& |
| YGPM5l18 | *[FMP30]*, YPL102C, ELP4, ATG21, FMP14, YPL098C, MSY1, ERI1, [PNG1]** |
| YGPM10h21 | *YPR158C-D, YPR158C-C, tA(AGC)P, KRE6, YPR159C-A, tG(GCC)P2, [GPH1]*, YPR160W-A, [YPR160C-A]* |

| | |
|---|---|
| YGPM9k24 | *[MMS1]&, RHO1, MRP2, MET16, NUT2, JIP5, tI(AAU)P2, YPR170C, YPR169W-A, YPR170W-B, YPR170W-A, BSP1, [YPR172W]** |

[ ]* indicates that the 3' end of the gene is missing.
[ ]& indicates that the 5' end of the gene is missing.
[ ] indicates that the ORF is intact, but might be missing necessary upstream or downstream sequences.

**Supplemental Table S3 -** Strains used in this study:

All strains are derivatives of W303 (*ade2-1 his3-11, 15 leu2-3,112 trp1-1 ura3-1 can1-100*) unless otherwise noted.

| Strain name | Genotype |
|---|---|
| LY4027 | *MATa/α, $P_{TUB1}$-GFP-TUB1:LEU2/ $P_{TUB1}$-GFP-TUB1:LEU2, ZIP1-GFP/+, SPC42-mCherry:kanMX/+* |
| LY4273 | *MATa/α, $P_{TUB1}$-GFP-TUB1:URA3/+, ZIP1-GFP/+, SPC42-mCherry:kanMX/+, ndt80::kanMX/ ndt80::kanMX, $P_{NDT80-mse1Δ, mse1Δ}$NDT80:HIS3/ $P_{NDT80-mse1Δ, mse1Δ}$NDT80:HIS3, ADE2/ADE2, TRP1/TRP1* |
| LY4404 | *MATa/α, $P_{TUB1}$-GFP-TUB1:URA3/+, ZIP1-GFP/+, SPC42-mCherry:kanMX/+, ndt80::kanMX/ ndt80::kanMX, $P_{NDT80-mse1Δ, mse1Δ}$NDT80:HIS3/ $P_{NDT80-mse1Δ, mse1Δ}$NDT80:HIS3, ADE2/ADE2, TRP1/TRP1, pLB227 (2μ)* |
| LY4285 | *MATa/α, $P_{TUB1}$-GFP-TUB1:URA3/+, ZIP1-GFP/+, SPC42-mCherry:kanMX/+, ndt80::kanMX/ ndt80::kanMX, $P_{NDT80-mse1Δ, mse1Δ}$NDT80:HIS3/ $P_{NDT80-mse1Δ, mse1Δ}$NDT80:HIS3, ADE2/ADE2, TRP1/TRP1, $P_{NDT80}$-NDT80:LEU2 (2μ)* |
| LY5463 | *MATa/α, $P_{TUB1}$-GFP-TUB1:URA3/+, ZIP1-GFP/+, SPC42-mCherry:kanMX/+, ndt80::kanMX/ ndt80::kanMX, $P_{NDT80-mse1Δ, mse1Δ}$NDT80:HIS3/ $P_{NDT80-mse1Δ, mse1Δ}$NDT80:HIS3, ADE2/ADE2, TRP1/TRP1, $P_{BCY1}$-BCY1:LEU2 (2μ)* |
| LY8715 | *MATa/α, $P_{TUB1}$-GFP-TUB1:LEU2/ $P_{TUB1}$-GFP-TUB1:LEU2, ZIP1-GFP/+, SPC42-mCherry:kanMX/+, bcy1::NAT/+* |
| LY5465 | *MATa/α, $P_{TUB1}$-GFP-TUB1:URA3/+, ZIP1-GFP/+, SPC42-mCherry:kanMX/+, ndt80::kanMX/ ndt80::kanMX, $P_{NDT80-mse1Δ, mse1Δ}$NDT80:HIS3/ $P_{NDT80-mse1Δ, mse1Δ}$NDT80:HIS3, ADE2/ADE2, TRP1/TRP1, $P_{IME2}$-IME2:LEU2 (2μ)* |
| LY8932 | *MATa/α, $P_{TUB1}$-GFP-TUB1:URA3/ $P_{TUB1}$-GFP-TUB1:URA3, ZIP1-GFP/+, SPC42-mCherry:HPH/+, ime2::HPH/ ime2::HPH, $P_{IME2}$-IME2-myc-TRP1:TRP1/ $P_{IME2}$-IME2-myc-TRP1:TRP1, sum1::kanMX/sum1::NAT* |
| LY5430 | *MATa/α, $P_{TUB1}$-GFP-TUB1:URA3/ $P_{TUB1}$-GFP-TUB1:URA3, ZIP1-GFP/+, SPC42-mCherry:kanMX/+, ime2::HPH/ ime2::HPH, $P_{IME2}$-IME2 T242A-myc-TRP1:TRP1/ $P_{IME2}$-IME2 T242A-myc-TRP1:TRP1* |
| LY5584 | *MATa/α, $P_{TUB1}$-GFP-TUB1:URA3/+, ZIP1-GFP/+, SPC42-mCherry:kanMX/+, ndt80::kanMX/ ndt80::kanMX, $P_{NDT80-mse1Δ, mse1Δ}$NDT80:HIS3/ $P_{NDT80-mse1Δ, mse1Δ}$NDT80:HIS3, ADE2/ADE2, TRP1/TRP1, $P_{BMH1}$-BMH1:LEU2 (2μ)* |
| LY8279 | *MATa/α, $P_{TUB1}$-GFP-TUB1:URA3/+, ZIP1-GFP/+, SPC42-mCherry:kanMX/+, ndt80::kanMX/ ndt80::kanMX, $P_{NDT80-mse1Δ, mse1Δ}$NDT80:HIS3/ $P_{NDT80-mse1Δ, mse1Δ}$NDT80:HIS3, ADE2/ADE2, TRP1/TRP1, $P_{BMH2}$-BMH2:LEU2 (2μ)* |
| LY5663 | *MATa/α, $P_{TUB1}$-GFP-TUB1:LEU2/ $P_{TUB1}$-GFP-TUB1:LEU2, ZIP1-GFP/+, SPC42-mCherry:kanMX/+, bmh1::NAT/bmh1::NAT* |

| | |
|---|---|
| LY5837 | *MATa/α, $P_{TUB1}$-GFP-TUB1:LEU2/ $P_{TUB1}$-GFP-TUB1:LEU2, ZIP1-GFP/+, SPC42-mCherry:kanMX/+, bmh2::NAT/bmh2::NAT* |
| LY6884 | *MATa/α, $P_{TUB1}$-GFP-TUB1:LEU2/ $P_{TUB1}$-GFP-TUB1:LEU2, ZIP1-GFP/+, SPC42-mCherry:hphMX/+, bmh2::NAT/bmh2::NAT, kanMX:$P_{CLB2}$-BMH1/ kanMX:$P_{CLB2}$-BMH1* |
| LY6948 | *MATa/α, $P_{TUB1}$-GFP-TUB1:LEU2/ $P_{TUB1}$-GFP-TUB1:LEU2, ZIP1-GFP/+, SPC42-mCherry:hphMX/+, bmh2::NAT/bmh2::NAT, kanMX:$P_{CLB2}$-BMH1/ kanMX:$P_{CLB2}$-BMH1, mek1::HIS3/mek1::HIS3* |
| LY6963 | *MATa/α, $P_{TUB1}$-GFP-TUB1:LEU2/ $P_{TUB1}$-GFP-TUB1:LEU2, ZIP1-GFP/+, SPC42-mCherry:hphMX/+, bmh2::NAT/bmh2::NAT, kanMX:$P_{CLB2}$-BMH1/ kanMX:$P_{CLB2}$-BMH1, spo11::HIS3/spo11::HIS3* |
| LY6981 | *MATa/α, $P_{TUB1}$-GFP-TUB1:LEU2/ $P_{TUB1}$-GFP-TUB1:LEU2, ZIP1-GFP/+, SPC42-mCherry:hphMX/+, bmh2::NAT/bmh2::NAT, kanMX:$P_{CLB2}$-BMH1/ kanMX:$P_{CLB2}$-BMH1, mec1::LEU2/mec1::LEU2, sml1Δ/sml1Δ* |
| LY6257 | *MATa/α, $P_{TUB1}$-GFP-TUB1:URA3/ $P_{TUB1}$-GFP-TUB1:URA3, ZIP1-GFP/+, SPC42-mCherry:kanMX/+, cdc20::$P_{CLB2}$-3HA-CDC20::kanMX6/ cdc20::$P_{CLB2}$-3HA-CDC20::kanMX6* |
| LY6250 | *MATa/α, $P_{TUB1}$-GFP-TUB1:URA3/ $P_{TUB1}$-GFP-TUB1:URA3, ZIP1-GFP/+, SPC42-mCherry:kanMX/+, cdc20::$P_{CLB2}$-3HA-CDC20::kanMX6/ cdc20::$P_{CLB2}$-3HA-CDC20::kanMX6, bmh1::HPH/bmh1::HPH* |
| LY6284 | *MATa/α, $P_{TUB1}$-GFP-TUB1:URA3/ $P_{TUB1}$-GFP-TUB1:URA3, ZIP1-GFP/+, SPC42-mCherry:kanMX/+, cdc20::$P_{CLB2}$-3HA-CDC20::kanMX6/ cdc20::$P_{CLB2}$-3HA-CDC20::kanMX6, bmh2::HPH/bmh2::HPH* |
| LY7903 | *MATa/α, $P_{TUB1}$-GFP-TUB1:LEU2/ $P_{TUB1}$-GFP-TUB1:LEU2, ZIP1-GFP/+, SPC42-mCherry:kanMX/+, bmh1::NAT/bmh1::NAT, $P_{NDT80}$-NDT80:URA3 (2μ)* |
| LY7905 | *MATa/α, $P_{TUB1}$-GFP-TUB1:LEU2/ $P_{TUB1}$-GFP-TUB1:LEU2, ZIP1-GFP/+, SPC42-mCherry:kanMX/+, bmh2::NAT/bmh2::NAT, $P_{NDT80}$-NDT80:URA3 (2μ)* |
| LY3273 | *MATa/α, $P_{TUB1}$-GFP-TUB1:URA3/ $P_{TUB1}$-GFP-TUB1:URA3, ZIP1-GFP/+, SPC42-mCherry:kanMX/+, cdc20::$P_{CLB2}$-3HA-CDC20::kanMX6/ cdc20::$P_{CLB2}$-3HA-CDC20::kanMX6* |
| LY2289 | *MATa/α, $P_{TUB1}$-GFP-TUB1:URA3/ $P_{TUB1}$-GFP-TUB1:URA3, ZIP1-GFP/+, SPC42-mCherry:kanMX/+, kanMX:$P_{CLB2}$-CDC5/ kanMX:$P_{CLB2}$-CDC5* |
| LY3515 | *MATa/α, $P_{TUB1}$-GFP-TUB1:URA3/ $P_{TUB1}$-GFP-TUB1:URA3, ZIP1-GFP/+, SPC42-mCherry:HPH/+, cdc5::NAT/+* |
| LY3518 | *MATa/α, $P_{TUB1}$-GFP-TUB1:URA3/ $P_{TUB1}$-GFP-TUB1:URA3, ZIP1-GFP/+, SPC42-mCherry:HPH/+, cdc5L158G:NAT/ cdc5L158G:NAT* |
| LY3639 | *MATa/α, $P_{TUB1}$-GFP-TUB1:URA3/ $P_{TUB1}$-GFP-TUB1:URA3, ZIP1-GFP/+, SPC42-mCherry:HPH/+, cdc5L158G:NAT/ cdc5L158G:NAT, cdc20::$P_{CLB2}$-3HA-CDC20::kanMX6/ cdc20::$P_{CLB2}$-3HA-CDC20::kanMX6* |

| | |
|---|---|
| LY8522 | SEY6210 *MATa, his3Δ200 trp1-90 leu2-3,112 ade2 gal80 lys2::*$lexA_{op}$*-HIS3::LYS2 ura3::* $lexA_{op}$*-lacZ::URA3* |
| LY8770 | SEY6210 *MATa, his3Δ200 trp1-90 leu2-3,112 ade2 gal80 lys2::*$lexA_{op}$*-HIS3::LYS2 ura3::* $lexA_{op}$*-lacZ::URA3, GAL4AD-NDT80(284-627):LEU2 (2µ), lexA:TRP1 (2µ)* |
| LY8772 | SEY6210 *MATa, his3Δ200 trp1-90 leu2-3,112 ade2 gal80 lys2::*$lexA_{op}$*-HIS3::LYS2 ura3::* $lexA_{op}$*-lacZ::URA3, GAL4AD-NDT80(284-627):LEU2 (2µ), lexA-BMH1:TRP1 (2µ)* |
| LY8767 | SEY6210 *MATa, his3Δ200 trp1-90 leu2-3,112 ade2 gal80 lys2::*$lexA_{op}$*-HIS3::LYS2 ura3::* $lexA_{op}$*-lacZ::URA3, GAL4AD:LEU2 (2µ), lexA:TRP1 (2µ)* |
| LY8768 | SEY6210 *MATa, his3Δ200 trp1-90 leu2-3,112 ade2 gal80 lys2::*$lexA_{op}$*-HIS3::LYS2 ura3::* $lexA_{op}$*-lacZ::URA3, GAL4AD-CDC5:LEU2 (2µ), lexA:TRP1 (2µ)* |
| LY8769 | SEY6210 *MATa, his3Δ200 trp1-90 leu2-3,112 ade2 gal80 lys2::*$lexA_{op}$*-HIS3::LYS2 ura3::* $lexA_{op}$*-lacZ::URA3, GAL4AD:LEU2 (2µ), lexA-BMH1:TRP1 (2µ)* |
| LY8771 | SEY6210 *MATa, his3Δ200 trp1-90 leu2-3,112 ade2 gal80 lys2::*$lexA_{op}$*-HIS3::LYS2 ura3::* $lexA_{op}$*-lacZ::URA3, GAL4AD-CDC5:LEU2 (2µ), lexA-BMH1:TRP1 (2µ)* |
| LY7771 | *MATa/α,* $P_{TUB1}$*-GFP-TUB1:LEU2/* $P_{TUB1}$*-GFP-TUB1:LEU2, ZIP1-GFP/+, SPC42-mCherry:kanMX/+, bmh1::NAT/bmh1::NAT,* $P_{CDC5}$*-CDC5:URA3 (2µ)* |
| LY7790 | *MATa/α,* $P_{TUB1}$*-GFP-TUB1:LEU2/* $P_{TUB1}$*-GFP-TUB1:LEU2, ZIP1-GFP/+, SPC42-mCherry:kanMX/+, bmh2::NAT/bmh2::NAT,* $P_{CDC5}$*-CDC5:URA3 (2µ)* |
| LY8780 | *MATa/α,* $P_{TUB1}$*-GFP-TUB1:LEU2/* $P_{TUB1}$*-GFP-TUB1:LEU2, ZIP1-GFP/+, SPC42-mCherry:kanMX/+, bmh1::NAT/bmh1::NAT,* $P_{BMH1}$*-BMH1 3A:HIS3/* $P_{BMH1}$*-BMH1 3A:HIS3* (at *BMH1* locus) |
| LY8592 | *MATa/α,* $P_{TUB1}$*-GFP-TUB1:LEU2/* $P_{TUB1}$*-GFP-TUB1:LEU2, ZIP1-GFP/+, SPC42-mCherry:kanMX/+, pes4::NAT/pes4::NAT* |
| LY8934 | *MATa/α,* $P_{TUB1}$*-GFP-TUB1:LEU2/* $P_{TUB1}$*-GFP-TUB1:LEU2, ZIP1-GFP/+, SPC42-mCherry:kanMX/+, bmh1::NAT/bmh1::NAT,* $P_{PES4}$*-PES4:URA3 (2µ)* |
| LY8489 | *MATa/α, SPC42-mCherry:HPH/+, BMH1-EGFP:HIS5/ BMH1-EGFP:HIS5* |
| LY2906 | *MATa/α, SPC42-mCherry:kanMX/+* |
| B1421 | SK1 *MATa ho::LYS2 lys2 ura3 leu2::hisG his3::hisG trp1::hisG* |
| B2424 | SK1 *MATa ho::LYS2 lys2 ura3 leu2::hisG his3::hisG trp1::hisG* <br> *CDC5-3V5::G418R* |
| B48 | SK1 *MATa/α ho::LYS2/ho::LYS2 lys2/lys2 ura3/ura3 leu2::hisG/leu2::hisG his3::hisG/his3::hisG trp1::hisG/trp1::hisG ura3::*$P_{GPD1}$- |

| | |
|---|---|
| | *GAL4(848).ER::URA3/ura3::*$P_{GPD1}$*-GAL4(848).ER::URA3* $P_{GAL}$*-NDT80::TRP1/*$P_{GAL}$*-NDT80::TRP1 CLB3-3HA:KanR/CLB3-3HA:KanR RIM4-3V5::HIS3/RIM4-3V5::HIS3* |

**Supplemental Table S4.** Oligos used in this study.

| Oligo number | Sequence | Purpose |
|---|---|---|
| LO1677 | 5’ AAGTCTGTCGACGAAGGTTAAACGTGGCATTGAC 3’ | *BMH1* cloning |
| LO1678 | 5’ TCGAATGGATCCGTTGCTTCTCGCTATACCAGAG 3’ | *BMH1* cloning |
| LO1968 | 5’ TTGTCAGTCGACCTTCCACAACCACCTTCATCTTCTC 3’ | *BMH2* cloning |
| LO1969 | 5’ GCTATACCCGGGCATCCTTTCGTTTGTGATCCCTTGTG 3’ | *BMH2* cloning |
| LO1894 | 5’ CAAAATTGAGAGCGCAAGCAAGTGAGAAGCGTACGC TGCAGGTCG 3’ | *BMH1* deletion |
| LO1895 | 5’ AATTGCAATAATGAACTACAAATTATTACACCCCCGTTC TCATCGATGAATTCGAGCTCG 3’ | *BMH1* deletion |
| LO1936 | 5’ GCAAAGAACGAGAAAACTTGGACAGAAGGTTAATACT CTGACAACGTACGCTGCAGGTCG 3’ | *BMH2* deletion |
| LO1937 | 5’ GTGGTAAATCTTCATTTCCCCTTGTATTTCTCAGCGC TCTCATCGATGAATTCGAGCTCG 3’ | *BMH2* deletion |
| LO2254 | 5’ AACACCGAAAACAAAATCGGCTTTTTACGTGACAA AAGAAGGTTCGTACGCTGCAGGTCG 3’ | Promoter replacement of *BMH1* |
| LO2256 | 5’ TCTTCACGACTGGTTGACATTTTTATCTTTAG TTCTATAAGATCAATGAAGAGAGAGAGG 3’ | Promoter replacement of *BMH1* |
| LO659 | 5’ CCTTTTTATTTTGAACTACGCGCAGTTTGAAAGAGGAA GAGATAATCCTATGATTAAATCGATGAATTCGAGCTCG 3’ | *MEK1* deletion |
| LO660 | 5’ CGTTTCAAATTTTATTAAGGACCAAATATACAACAGAAA GAAGAAGAGCGGAATGCGTACGCTGCAGGTCGAC 3’ | *MEK1* deletion |
| LO64 | 5’ CTTCACCCTTAAGATTTTACGATTTACTAAGTTCACCTTCTC ATGCGTACGCTGCAGGTCG 3’ | *SPO11* deletion |
| LO65 | 5’ CTTGAAAAACATTTTTTATAAAGCAACAGCTCCC ATTCTTATTCAGATATCATCGATGAATTCGAGCTCG 3’ | *SPO11* deletion |
| LO2800 | 5’ CGTATGGGATCCTCAAGCCTATCCTCTTTCATCTCAGC 3’ | *CDC5* cloning |
| LO2801 | 5’ TTGTCAGAGCTCGATGAGAAGTTCTGGCTACCTTAGTC 3’ | *CDC5* cloning |

| | | |
|---|---|---|
| LO3062 | 5’ acgctagcttgggtggtcatatggccATGTCGTTGGGTCCTCTTAA 3’ | *CDC5* cloning in Y2H plasmid |
| LO3063 | 5’ tcagtatctacgattcatagatctcGATGAGAAGTTCTGGCT ACCTTAGTC 3’ | *CDC5* cloning in Y2H plasmid |
| LO3120 | 5’ AATCAGTATCACAAAAAAAATTTCAACCTACAAGAGG AAACATGCGTACGCTGCAGGTCG 3’ | *PES4* deletion |
| LO3121 | 5’ ATAAGGAAAACTCTAGCTAAATTAACAAAGAGAC ATTCATCATCGATGAATTCGAGCTCG 3’ | *PES4* deletion |
| LO3163 | 5’ gtatctatgtcgacGGCGTGCGGCAGTATAAGTTC 3’ | *PES4* cloning |
| LO3164 | 5’ CTATGGCTgagctcGACGCGGGAATGGAAGAGATAGG 3’ | *PES4* cloning |
| LO1808 | 5’ TATGCAGTCGACAGGACAAAGAAGGCAAAGAACC 3’ | *BCY1* cloning |
| LO1809 | 5’ GTCAATGGATCCGGTGATTGTAGAGTTGGAGAGAGT 3’ | *BCY1* cloning |
| LO3153 | 5’ ACAAGCAGATTATTTTCAAAAGACAACAGTAAGAA TAAACGATGCGTACGCTGCAGGTCG 3’ | *BCY1* deletion |
| LO3154 | 5’ AATTCATGTGGATTTAAGATCGCTTCCCCTTTTT ACTTATCATCGATGAATTCGAGCTCG 3’ | *BCY1* deletion |
| LO1673 | 5’ TACATCGTCGACCTCTACCATGCATTTCGAACATGT 3’ | *IME2* cloning |
| LO1674 | 5’ ACGTTACCCGGGGTAGCCGCCAAGAGCAAGAAA 3’ | *IME2* cloning |
| LO2976 | 5’ AAAGTTTCATACATAATTAACAAAATTCGTTTGTTGCGGG GATG CGTACGCTGCAGGTCG 3’ | *SUM1* deletion |
| LO2977 | 5’ ATCTATTCTCGAAACTGCCCCAACGTACGGACCAGCTTA TCATCGATGAATTCGAGCTCG 3’ | *SUM1* deletion |
| LO2026 | 5’ GTAATTTCTCTTTAGATTTATCAGAATACTTATCATCGAT GAATTCGAGCTCG 3’ | Tagging *BMH1* C-terminus with EGFP |
| LO2027 | 5’ ACAGCAGCCACCTGCTGCCGCCGAAGGTGAAG CACCAAAGCGTACGCTGCAGGTCG 3’ | Tagging *BMH1* C-terminus with EGFP |
| LO3089 | 5’ GGATCGTCgaattcATGTCAACCAGTCGTGAAGA 3’ | *BMH1* cloning in Y2H plasmid |
| LO3090 | 5’ ATCTGGATGTCGACGGTGGAGGAATCAGAAGAGAGA 3’ | *BMH1* cloning in Y2H plasmid |

| | | |
|---|---|---|
| LO686 | 5’ GCATctcgagCAAAGGGAACGAAACCCAG 3’ | *IME2* cloning |
| LO687 | 5’ ATGCgcggccgcCAGTAGCCGCCAAGAGCAAG 3’ | *IME2* cloning |
| LO1627 | 5’ TTTCAACATGGCGTGCCAAACCAAAATC 3’ | *IME2* site-directed mutagenesis |
| LO1629 | 5’ ATAAAAACCCGTATgCGGCCTACGTTTCC 3’ | *IME2* site-directed mutagenesis |

**Supplemental Table S5.** Plasmids used in this study.

| Plasmid number | Genotype/Purpose | Reference |
|---|---|---|
| YEplac195 | *URA3 (2µ)* | Gietz, Sugino 1988 |
| YEplac181 | *LEU2 (2µ)* | Gietz, Sugino 1988 |
| pBTM116 | *lexA:TRP1 (2µ)* | Hollingsworth 2018 |
| pACTII | *GAL4AD:LEU2 (2µ)* | Hollingsworth 2018 |
| pLB227 | Empty vector for the screen | This study |
| pLB225 | $P_{NDT80}$*-NDT80:LEU2 (2µ)* | This study |
| pLB262 | $P_{BMH1}$*-BMH1:LEU2 (2µ)* | This study |
| pLB539 | $P_{BMH2}$*-BMH2:LEU2 (2µ)* | This study |
| pLB518 | *lexA-BMH1:TRP1 (2µ)* | This study |
| pLB491 | *GAL4AD-CDC5:LEU2 (2µ)* | This study |
| pLB463 | $P_{CDC5}$*-CDC5:LEU2 (2µ)* | This study |
| pLB465 | $P_{CDC5}$*-CDC5:URA3 (2µ)* | This study |
| pLB509 | $P_{BMH1}$*-BMH1:HIS3* | This study |
| pLB519 | $P_{BMH1}$*-BMH1 3A:HIS3* | This study |
| pLB513 | $P_{PES4}$*-PES4:URA3 (2µ)* | This study |
| pLB306 | $P_{BCY1}$*-BCY1:LEU2 (2µ)* | This study |
| pLB258 | $P_{IME2}$*-IME2:LEU2 (2µ)* | This study |
| pLB268 | $P_{IME2}$*-IME2-myc-TRP1* | This study |
| pLB285 | $P_{IME2}$*-IME2 T242A-myc-TRP1* | This study |
| pLB107 | $P_{NDT80}$*-NDT80:HIS3* | This study |
